# DNA replication-mediated error correction of ectopic CENP-A deposition maintains centromere identity

**DOI:** 10.1101/428557

**Authors:** Yael Nechemia-Arbely, Karen H. Miga, Ofer Shoshani, Aaron Aslanian, Moira A. McMahon, Ah Young Lee, Daniele Fachinetti, John R. Yates, Bing Ren, Don W. Cleveland

## Abstract

Chromatin assembled with the histone H3 variant CENP-A is the heritable epigenetic determinant of human centromere identity. Using genome-wide mapping and reference models for 23 human centromeres, CENP-A is shown in early G1 to be assembled into nucleosomes within megabase, repetitive α-satellite DNAs at each centromere and onto 11,390 transcriptionally active sites on the chromosome arms. Here we identify that DNA replication acts as an error correction mechanism to sustain centromere identity through the removal of the sites of CENP-A loading on the chromosome arms, while maintaining centromere-bound CENP-A with the same DNA sequence preferences as in its initial loading.

## Introduction

Correct chromosome segregation during mitosis is crucial to ensure each daughter cell will receive a complete set of chromosomes. This process relies on a unique chromatin domain known as the centromere. Human centromeres are located on megabase long ^1^ chromosomal regions and are comprised of tandemly repeated arrays of a ~171 bp element, termed α-satellite DNA ^2–4^ CENP-A is a histone H3 variant ^5, 6^ that replaces histone H3 in ~3% of α-satellite DNA repeats ^7, 8^, is flanked by pericentric heterochromatin containing H3K9me2/3 ^9^, and apparently spans on 1/3^rd^ to one half of α-satellite arrays in the centromeres of chromosomes X and Y ^10^. Despite the correlation between centromere location and the presence of α-satellite DNA repeats, α-satellite DNA sequences are neither sufficient nor essential for centromere identity ^2, 11, 12^. This has been demonstrated by several measures including identification of multiple examples ^13^ of acquisition of a new centromere (referred to as a neocentromere) at a new location coupled with inactivation of the original centromere on the same chromosome. Indeed, while α-satellite arrays incorporated into human artificial chromosomes (HACs) can nucleate active centromeres ^14–21^, they do so at low (5–8%) efficiencies.

All of this has led to a consensus view that mammalian centromeres are defined by an epigenetic mark ^2^. Use of gene replacement in human cells and fission yeast has identified the mark to be CENP-A-containing chromatin ^22^, which maintains and propagates centromere function indefinitely by recruiting CENP-C and the <u>c</u>onstitutive <u>c</u>entromere <u>a</u>ssociated <u>c</u>omplex (CCAN) ^23–26^. We ^8^ and others ^27^ have shown that the overwhelming majority of human CENP-A chromatin particles contain two molecules of CENP-A at all cell cycle points, with CENP-A chromatin bound at authentic centromeres protecting 133 bp of centromeric α-satellite-containing DNA from nuclease digestion ^8, 28^ before and after DNA replication ^8^. This evidence is consistent with an octameric nucleosome with DNA unwinding at all cell cycle points, and with no evidence for oscillation between hemisomes and octasomes, and with heterotypic CENP-A/histone H3-containing nucleosomes comprising at most 2% of CENP-A-containing chromatin ^8^.

During DNA replication, initially bound CENP-A is quantitatively redistributed to each daughter centromere ^29^, while incorporation of new molecules of CENP-A into chromatin occurs only for a short period after exit from mitosis ^29–32^ when its loading chaperone HJURP ^33, 34^ is active ^35^. This temporal separation of new CENP-A chromatin assembly at mitotic exit from centromeric DNA replication raises the important question of how is the epigenetic mark that determines centromere identity maintained across the cell cycle when it is expected to be dislodged by DNA replication and diluted at each centromere as no new CENP-A is assembled until the next G1 ^29^. Moreover, endogenous CENP-A comprises only ~0.1% of the total histone H3 variants. Recognizing that a proportion of CENP-A is assembled at the centromeres with the remainder loaded onto sites on the chromosome arms ^7, 8, 36^, long-term maintenance of centromere identity and function requires limiting accumulation of non-centromeric CENP-A. Indeed, artificially increasing CENP-A expression by several fold in human cells ^36–39^ or flies ^40^ or the CENP-A homolog (Cse4) in yeast ^41, 42^ increases ectopic deposition at non-centromeric sites, accompanied by chromosome segregation aberrations.

Using centromere reference models for each of the centromeres of the 22 human autosomes and the X chromosome, we show that after DNA replication centromere-bound CENP-A is reassembled into nucleosomes onto α-satellite DNA sequences with sequence preferences that are indistinguishable from those bound in its initial HJURP-dependent loading at mitotic exit and that this re-loading is independent of CENP-A expression level. Furthermore, we identify that a DNA synthesis-mediated error correction mechanism acts in S phase to remove ectopically loaded CENP-A found within transcriptionally active chromatin outside of the centromeres while retaining centromere-bound CENP-A, resulting in maintenance of epigenetically defined centromere identity.

## Results

### CENP-A binding at 23 human centromere reference models

To identify the sequences bound by CENP-A across each human centromere, chromatin was isolated from synchronized HeLa cells expressing either i) CENP-A^LAP^, a CENP-A variant carboxy-terminally fused to a localization [EYFP] and <u>a</u>ffinity [His] <u>p</u>urification tag ^43^ at one endogenous CENP-A allele (Fig. S1a, Fig. 1a) or ii) stably expressing an elevated level (4.5 times the level of CENP-A in parental cells) of CENP-A^TAP^, a CENP-A fusion with carboxy-terminal <u>t</u>andem <u>a</u>ffinity <u>p</u>urification (S protein and protein A) tags separated by a TEV protease cleavage site (Fig. S1b, Fig. 1a). Centromeric localization of both CENP-A variants was confirmed using immunofluorescence (Fig. S1c, d), each of which has previously been shown to support long-term centromere identity and mediate high fidelity chromosome segregation in the absence of wild type CENP-A ^7^, ^8^.

**Figure 1.**
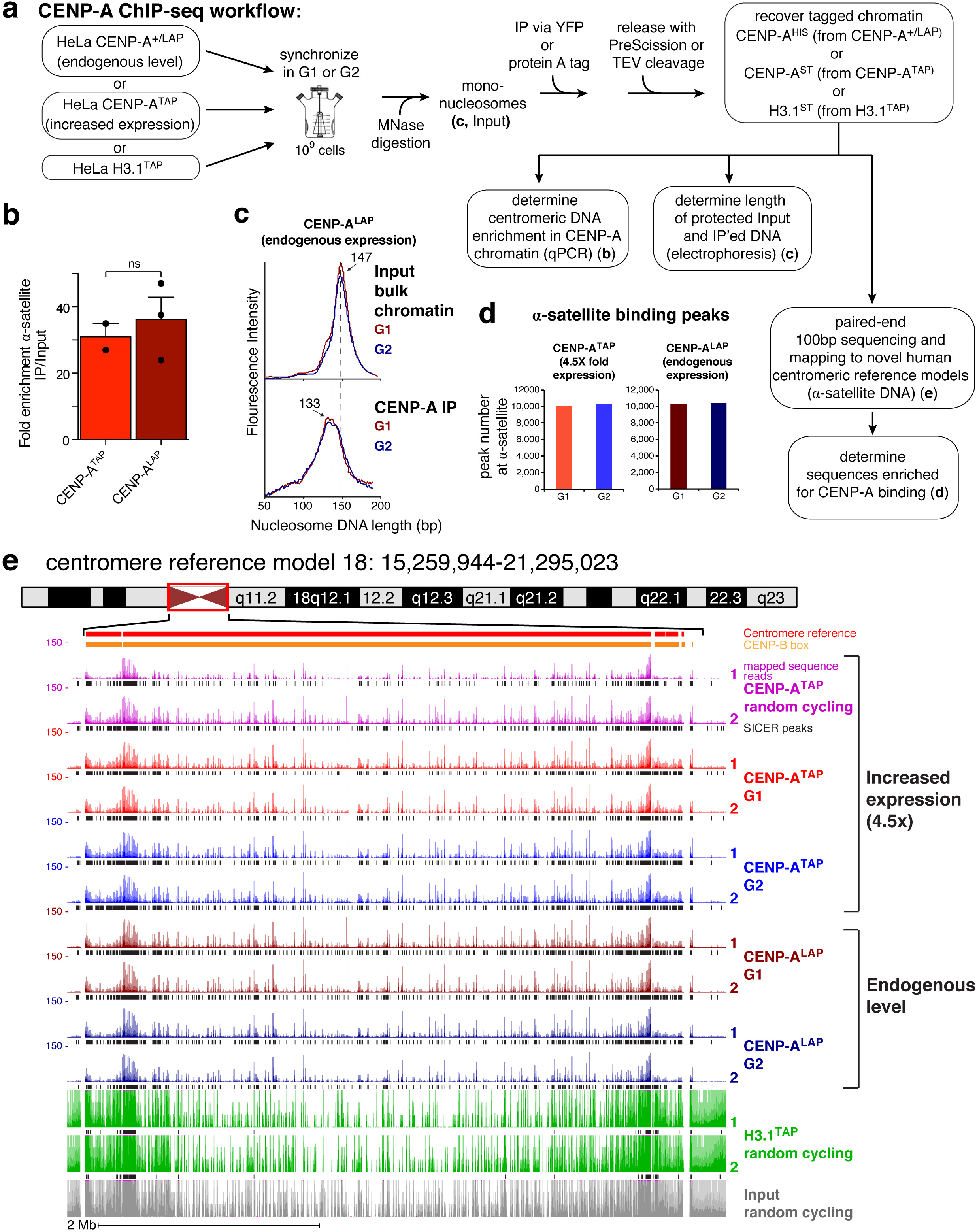
CENP-A ChIP-seq identifies CENP-A binding at reference centromeres of 23 human chromosomes. (**a**) CENP-A ChIP-sequencing experimental design. (**b**) Quantitative real-time PCR for α-satellite DNA in chromosomes 1, 3, 5, 10, 12 and 16. N=2 for CENP-A^TAP^ and 3 for CENP-A^LAP^, from two independent replicates. Error bars, s.e.m. *P* value of 0.608 determined using two-tailed *t*-test. (**c**) MNase digestion profile showing the nucleosomal DNA length distributions of bulk input mono-nucleosomes (upper panel) and purified CENP-A^LAP^ following native ChIP at G1 and G2. (**d**) Number of CENP-A binding peaks at α-satellite DNA in CENP-A^TAP^ and CENP-A^+/LAP^ cells at G1 and G2. The number represent peaks that are overlapping between the two replicates. (**e**) CENP-A ChIP-seq shows CENP-A binding peaks at the centromere of chromosome 18 for CENP-A^LAP^ and CENP-A^TAP^ before and after DNA replication. CENP-A peaks across the reference model are a result of multi-mapping and their exact linear order is not known. SICER peaks are shown in black below the raw read data. Two replicates are shown for each condition. Scale bar, 2Mb.

Chromatin was isolated from cells synchronized to be in G1 or G2 (Fig. 1a). Synchronization efficiency was >80% as determined by fluorescent activated cell sorting (FACS) analysis of DNA content (Fig. S1e). In parallel, chromatin was also isolated from randomly cycling cells stably expressing TAP tagged histone H3.1 (H3.1^TAP^ - Fig. S1b, Fig. 1a) ^23^. Chromatin from each line at G1 and G2 cell cycle phases was digested with micrococcal nuclease to generate mono-nucleosomes, producing the expected 147 bp of protected DNA length for bulk nucleosomes assembled with histone H3 (Fig. 1a, c - upper panel). CENP-A^LAP^, CENP-A^TAP^ or H3.1^TAP^ containing complexes were then affinity purified from the mono-nucleosome-containing pool using anti-GFP or rabbit-IgG antibodies coupled to magnetic beads. PreScission or TEV protease cleavage was then used to elute His or S-protein tagged CENP-A or S-tagged H3.1 chromatin under mild conditions that maintain initially assembled chromatin (Fig. 1a). α-satellite DNA sequences were enriched 30–35 fold in DNA isolated from CENP-A^TAP^ or CENP-A^+/LAP^ cells (Fig. 1b), the expected enrichment since α-satellite DNA comprises ~3% of the genome ^8, 21^. While micro-capillary electrophoresis of bulk input chromatin produced the expected 147 bp of protected DNA length for nucleosomes assembled with histone H3 (Fig. 1 c - upper panel), isolated CENP-A^LAP^ chromatin expressed at endogenous CENP-A levels produced DNA lengths centered on 133 bp, both before and after DNA replication (Fig. 1c, lower panel), a distribution indistinguishable from that previously reported for octameric CENP-A-containing nucleosomes assembled in vitro and in which DNA unwinding at entry and exit has been demonstrated ^8, 44^.

Libraries of affinity purified CENP-A^LAP^, CENP-A^TAP^, and H3.1^TAP^-bound DNAs were prepared, sequenced (using paired-end 100 bp sequencing), and mapped (Fig. 1a, d and Table S1) onto the published reference model for the centromere of the X chromosome ^45^ and unpublished reference models for the centromeres of the 22 human autosomes ^46, 47^. These centromere models include the observed variation in α-satellite <u>H</u>igher <u>O</u>rder <u>R</u>epeat (HOR) array sequences contained in the HuRef genome ^48^. The highly repeated sequences preclude distinguishing between centromeric and pericentromeric sequences and the order of repeats in the models is arbitrarily assigned and portions of the centromeres of the acrocentric chromosomes 13, 14, 21 and 22, as well as portions of centromeres of chromosomes 1, 5 and 19, contain nearly identical arrays that cannot be distinguished.

Sequences in each reference centromere associated with CENP-A binding were identified (Fig. 1d, e; Fig. S2) using algorithm-based scripts [SICER and MACS ^49, 50^]. Mapping of CENP-A^LAP^ expressed at endogenous levels across the centromeric regions of all 23 reference centromeres (see Fig. 1e for chromosome 18 and Figure S2 for the other 22) revealed the profile of CENP- A binding. Mapping to the sequences in the reference centromeres was highly reproducible (e.g., compare duplicates in Figs. 1e) and largely unaffected by increasing CENP-A levels by 4.5-fold (e.g., compare sequences bound in CENP-A^TAP^ cells with those in CENP-A+^/LAP^ cells in Figs. 1e and 2a, b).

**Figure 2.**
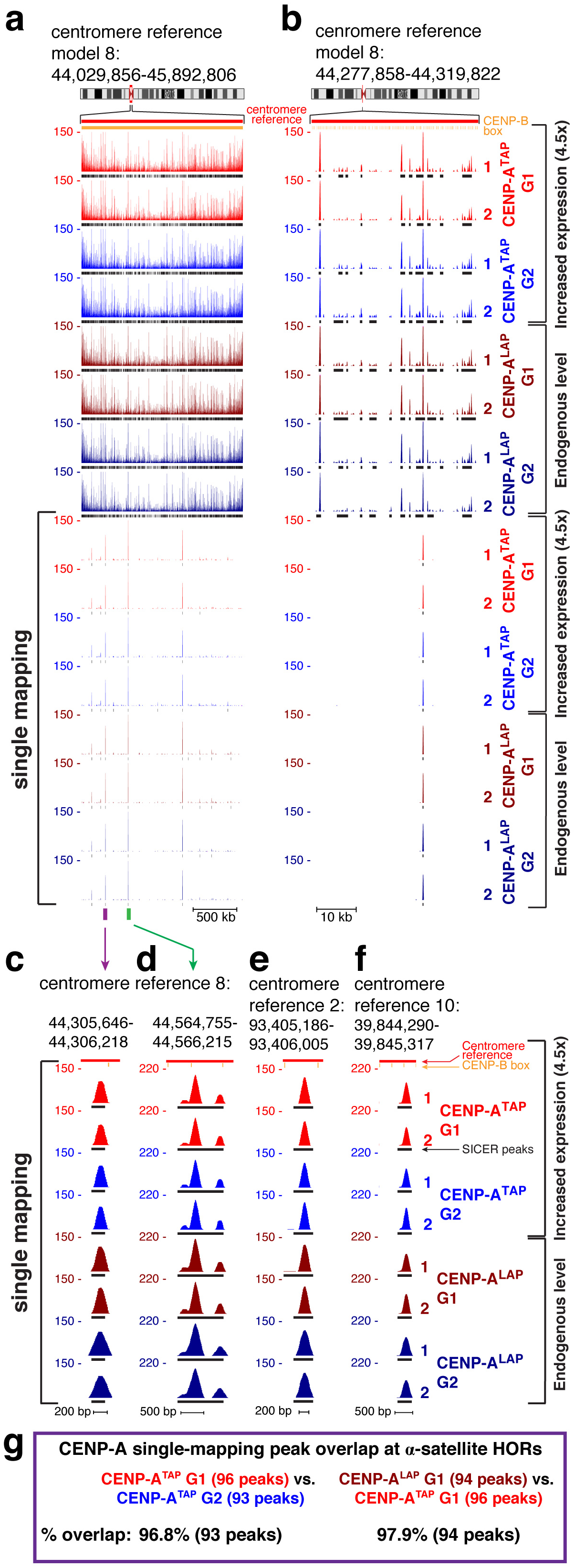
Retention of centromeric CENP-A through DNA replication. (**a, b**) CENP-A ChIP-seq raw mapping data (colored) and SICER peaks (black lines, underneath) showing sequences bound by CENP-A (at both endogenous and increased expression levels) across centromere reference model of chromosome 8, before and after DNA replication. Upper part shows mapping of all reads (including reads that are multi-mapping) onto the repetitive α−satellite DNA. Lower part shows read mapping to sites that are single copy in the HuRef genome (single-mapping), after filtering out multi-mapping reads. Centromere reference location, red. CENP-B box location, orange. Scale bar, 500kb (**a**), 10kb (**b**). (**c**) High-resolution view of read mapping to a site that is single copy in centromere reference model of chromosome 8, marked by a purple bar in (**a**). Scale bar, 200bp. (**d**) High-resolution view of read mapping to a site that is single copy in centromere reference model of chromosome 8, marked by a green bar in (**a**). Scale bar, 500bp. (**e, f**) High-resolution view of read mapping to a site that is single copy in centromere reference model of chromosome 2 (**e**) and in the centromere reference model of chromosome 10 (**f**). Scale bar, 200bp (**e**) and 500bp (**f**). (**g**) Left, overlap between G1 and G2 CENP-A single mapping binding sites at α-satellite HOR sequences. Right, peak overlap between G1 CENP-A^TAP^ (increased expression) and CENP-A^LAP^ (endogenous level) single mapping binding sites at α-satellite HOR sequences.

Analysis of CENP-A-bound DNAs aligned to α-satellite sequences in CENP-A+^/LAP^ G1 cells revealed that our CENP-A ChIP-seq approach resulted in varying levels of array enrichment, from ~10.5-fold enrichment for array D3Z1 in cen3 to ~213-fold for array GJ211930.1 in cen10, with most active arrays showing enrichment level of 20–40-fold above background (input) level (Table S2; columns 13, 14). For the 6 of the 17 centromeres which contain more than one α-satellite array within them, CENP-A was enriched only in one, indicating that only one array was active (see centromere reference models 3, 7, 12, 15, 16 and 19; Table S2: columns 13, 14). Multiple α-satellite arrays in 11 centromeres (for example, see centromere reference models 10,11,13, 14, and 17; Table S2: columns 13, 14) showed enrichment of CENP-A binding in two or more arrays. These may represent functional epialleles for CENP-A binding, (i.e., in which CENP-A binds to a different array in each homologue), as was shown previously for cen17 in two diploid cell human lines ^20^. Increased levels of CENP-A expression (in CENP-A^TAP^ versus CENP-A^+/LAP^ cells) did not increase the number of binding peaks in the reference centromeres (Fig. 1d, e), but did increase occupancy within the cell population at some divergent monomeric α-satellite repeats (Fig. S1f) or within HORs (Fig. S1g), with both examples occurring in regions with few 17 bp CENP-B boxes (Fig. S1f, g - compare sites bound at endogenous or increased CENP-A levels) that are direct binding sites for CENP-B ^51^, the only known sequence-specific human centromere binding protein.

### CENP-A nucleosomes are retained at centromeric loading sites after DNA replication

Next, we examined how centromeric CENP-A binding was affected by DNA replication. Despite the known redistribution of initially centromere-bound CENP-A onto each of the new daughter centromeres without addition of new CENP-A ^29^, comparison of the sequences bound by CENP- A in G1 with those bound in G2 revealed the remarkable feature that for all 23 centromeres, at both normal (CENP-A+^/LAP^) and elevated (CENP-A^TAP^) levels, CENP-A was bound to indistinguishable α-satellite sequences before and after DNA replication (shown for the reference centromere of chromosome 18 in Fig. 1e and for the other chromosomes in Fig. S2). Indeed, most (87%) of α-satellite sequences with CENP-A binding peaks in chromatin immunopurified from G1 CENP-A^TAP^ cells remained at G2 (Fig. S1h, top). Similarly high overlap (89%) was identified between repeats (called by SICER ^50^) bound by CENP-A in G1 or G2 when CENP-A was expressed at endogenous levels (in CENP-A+^/LAP^ cells) (Fig. S1h, bottom).

To address whether CENP-A was precisely retained at the same centromeric α-satellite loading sites after DNA replication, we filtered out multi-mapping reads, leaving only reads that map to sites that are single copy in the HuRef genome, and therefore are mapped uniquely. Analysis of these single copy variants revealed that CENP-A was quantitatively retained after DNA replication at single copy sites found within HORs (Fig. 2a-f). Indeed, almost all (93 of 96) unique CENP-A binding sites mapped within the 23 centromeres of CENP-A^TAP^ cells in G1 remained undiminished at G2 (Fig. 2g, left), with the remaining 3 peaks only slightly diminished.

### Sites of CENP-A assembly onto chromosome arms in early G1 are removed by G2

Genome-wide mapping of CENP-A-bound DNAs revealed that, in addition to the striking enrichment at centromeric α-satellites, CENP-A was preferentially and highly reproducibly incorporated into unique sequence, non-α-satellite sites on the arms of all 23 chromosomes (Figs. 3a, b and 4). Sites enriched for CENP-A binding were essentially identical in DNAs from randomly cycling cells or G1 cells (Fig. 3a, b, mapped sequence reads in color and binding sites underneath in black). A 4.5-fold increase in CENP-A levels in CENP-A^TAP^ cells drove correspondingly increased CENP-A sites of incorporation on the arms (from 11,390 sites at normal CENP-A levels to 40,279 sites when CENP-A was elevated 4.5-fold - Fig. 3a-b, d), but did not increase the binding peaks within the centromeric HORs (Fig. 1d) or the number of unique single copy sites within centromeric HORs (Fig. 2g, right).

**Figure 3.**
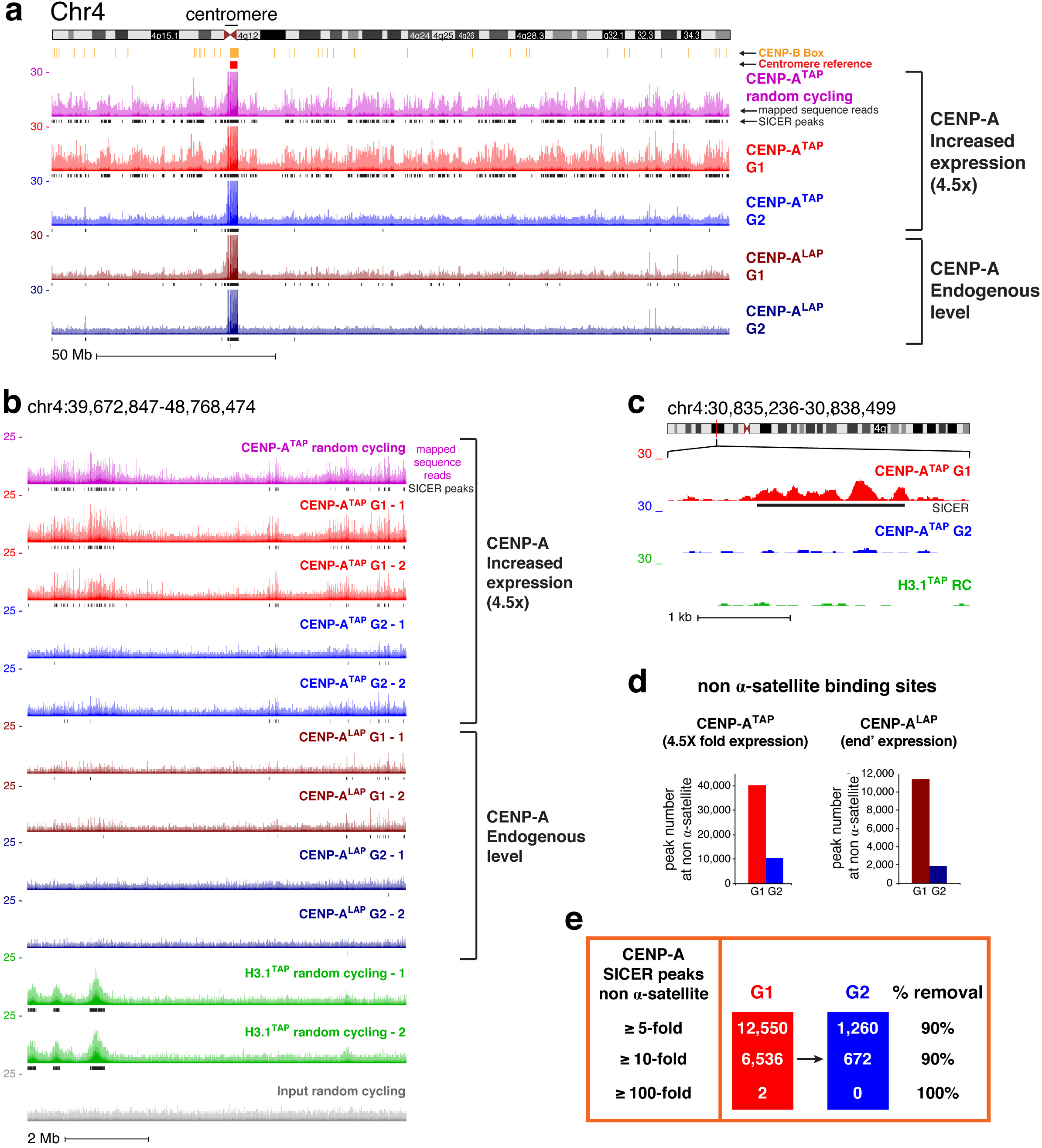
Sites of CENP-A assembly onto chromosome arms in early G1 are removed by G2. (**a**) ChIP-sequencing raw mapping data (colored) and SICER peaks (black lines, underneath) showing sequences bound by CENP-A (at both endogenous and increased expression levels) across chromosome 4 before and after DNA replication. Centromere reference location, red. CENP-B box location, orange. Read counts were scale to 30 but reaches 150 at the centromere. Scale bar, 50Mb. (**b**) ChIP-sequencing data are shown for a region within the p-arm of chromosome 4, with two replicates for each time point, for CENP-A^TAP^ (increased CENP-A expression), CENP-A^LAP^ (endogenous level), and H3.1^TAP^. Scale bar, 2Mb. (**c**) High resolution nucleosomal view of CENP-A^TAP^ mapping data at G1 and G2 at a non-centromeric site of chromosome 4. Scale bar, 1kb. (**d**) Total number of non-α-satellite CENP-A binding sites for CENP-A^TAP^ and CENP-A^LAP^ at G1 and G2. The number represent peaks that are overlapping between the two replicates. (**e**) Number of non-α-satellite CENP-A SICER binding sites called at G1 or G2 at different fold thresholds (above background).

Remarkably, for all 23 human chromosomes and for CENP-A accumulated to endogenous (CENP-A^LAP^) or increased (CENP-A^TAP^) expression levels, passage from G1 to G2 almost eliminated enrichment of CENP-A binding to specific sites on the chromosome arms, while leaving α-satellite bound sequences unaffected (Figs. 1d, 3, 4). Loss by G2 of CENP-A binding in G1 at specific arm sites was highly reproducible, as demonstrated by experimental replicas (Fig. 3b). On a genome-wide scale, scoring peak binding sites with thresholds of ≥ 5-fold, 10-fold or 100-fold of CENP-A binding over background, at least 90% of sites bound on chromosome arms in G1 in CENP-A^TAP^ cells were removed by early G2 (Fig. 3e) and all of those still identified in G2 were substantially reduced in peak height.

**Figure 4.**
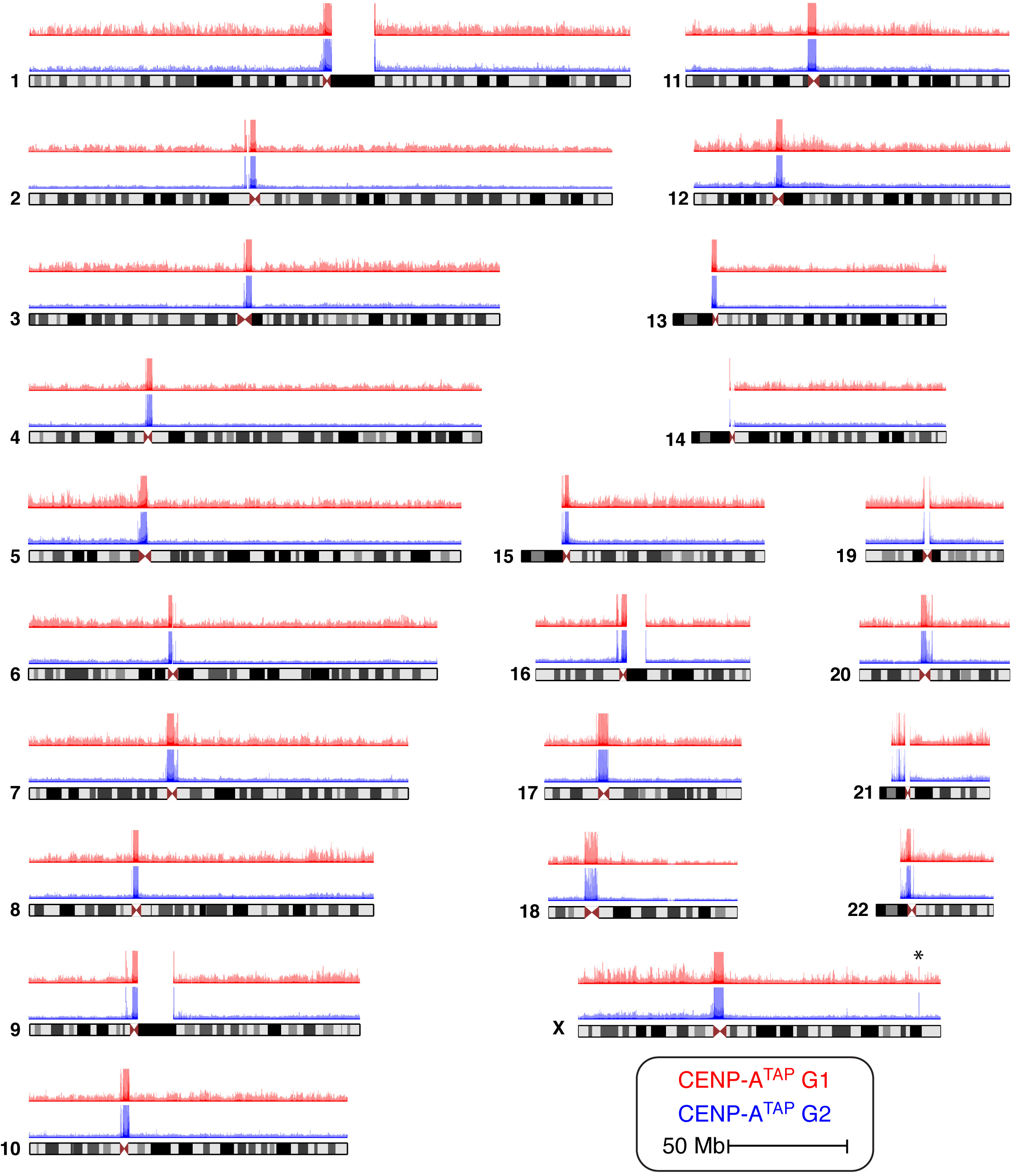
Ectopic CENP-A is removed following DNA replication from the arms of all 23 human chromosomes. CENP-A^TAP^ ChIP-sequencing raw mapping data at G1 (red) and G2 (blue) for all human chromosomes. Chromosome X show a spike of CENP-A enrichment not removed by G2 (marked by an asterisk).

Taken together, whether at endogenous or increased CENP-A expression level, CENP-A is loaded in G1 not only at centromeric α-satellite DNAs but also at preferential sites on the chromosome arms, but passage across S phase removes or sharply diminishes these unique sequence sites of enhanced CENP-A binding on the arms, while CENP-A binding to centromeric sequences is retained.

### CENP-A loading sites on chromosome arms are not seeding hotspots for neocentromere formation

We next determined whether sites of preferential CENP-A loading onto the chromosome arms corresponded to sites that have become active “neocentromere” locations ^52^ for the three human neocentromeres for which prior work has defined their chromosomal locations ^28^. The first of these, named PDNC4, has a neocentromere on chromosome 4 ^53^ that spans 300 kb (from 88.2 to 88.5 Mb). No peak binding sites of CENP-A relative to neighboring regions were found in this chromosomal region even with elevated CENP-A expression in CENP-A^TAP^ cells (Fig. 5a). Similar examination of two additional neocentromere-containing locations/positions [line MS4221 that harbors a 400 kb neocentromere at position 86.5 to 86.9 Mb on chromosome 8 ^28, 54^ and line IMS13q with a neocentromere on chromosome 13 ^55^ that spans 100 kb (from 97.7 to 97.8 Mb)] again revealed no elevated incorporation of CENP-A^TAP^ within the DNA sequences corresponding to the neocentromere domains (Fig. S3a, b).

**Figure 5.**
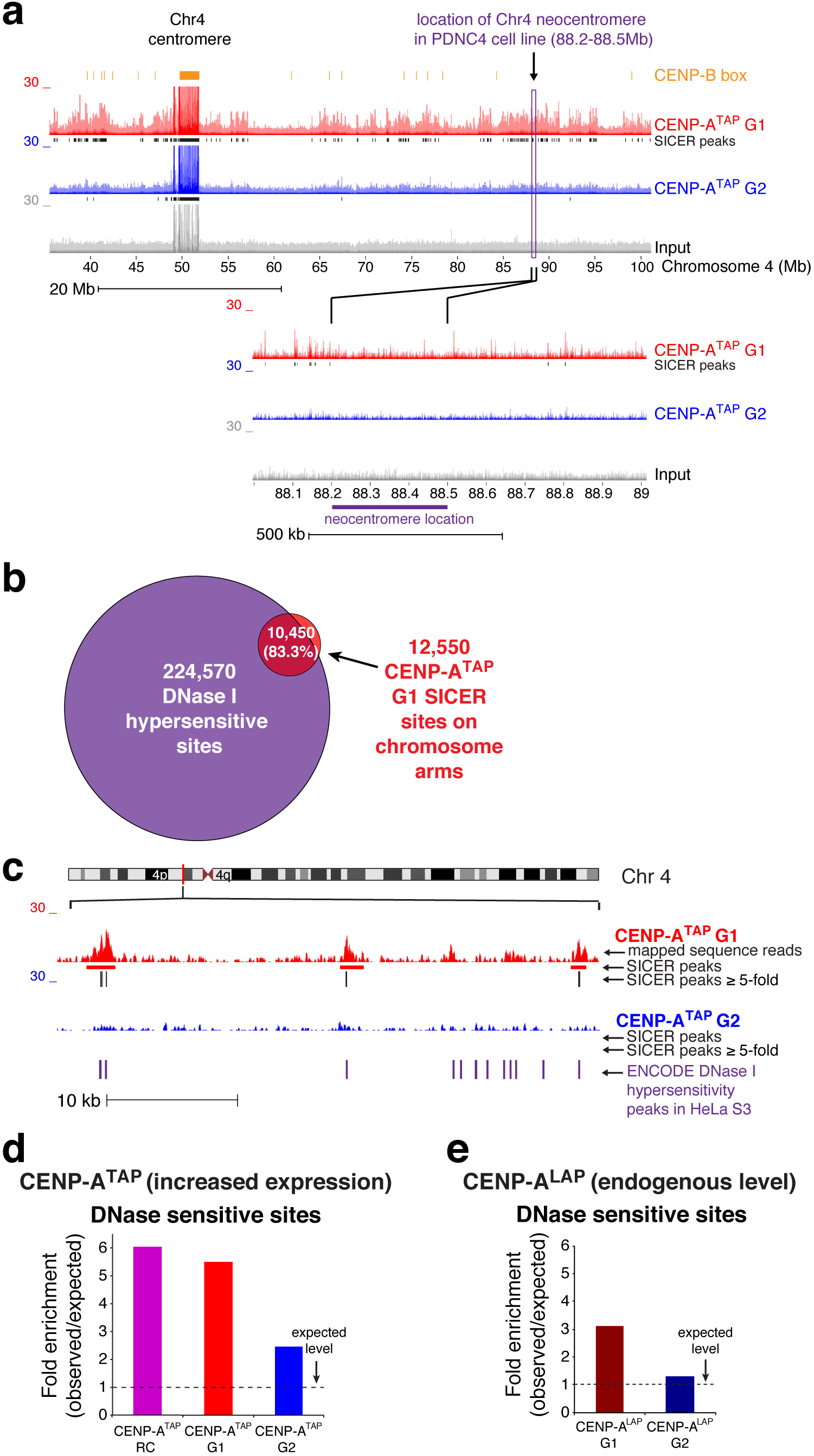
Sites of deposition of CENP-A on chromosome arms are not seeding hotspots for neocentromere formation. (**a**) Read mapping data of CENP-A^TAP^ ChIP-sequencing at G1 and G2, at the chromosomal location of a known patient derived neocentromere ^28^ found in chromosome 4. (**b**) Greater than 80% of CENP-A SICER peaks ≥ 5-fold in randomly cycling and G1 cells overlap with DNase I hypersensitive sites taken from ENCODE project. (**c**) Example from the chromosome 4 p-arm showing overlap of at least 100 bases between SICER peaks ≥5-fold and HeLa S3 DNase I hypersensitive sites taken from ENCODE project. (**d, e**) CENP-A^TAP^ (d) or CENP-A^LAP^ (e) enrichement levels at DNase I hypersensitive sites. SICER peaks ≥5-fold supported between the two replicates were analyzed for their enrichment level at DNase I hypersensitive sites, with minimum overlap of 100 bases, compared to the level of enrichment at these sites by chance.

### CENP-A is ectopically loaded at early G1 into open and active chromatin

We examined the nature of the sites on the chromosome arms into which CENP-A was assembled in G1. A 2-fold enrichment (compared to levels expected by chance) of CENP-A^TAP^ bound to unique arms sites during G1 was found at promoters, enhancers or promoters of expressed genes and a 2.5-fold enrichment at sites bound by the transcriptional repressor CTCF (Fig. S3d), a similar trend to what has been observed previously for increased expression of CENP-A ^36^. Importantly, more than 80% of CENP-A^TAP^ binding sites on chromosome arms with peak heights ≥ 5-fold over background overlapped with DNase I hypersensitive sites identified by comparison with ENCODE DNase I hypersensitive datasets (with minimum overlap of 100 bases) that denote accessible chromatin zones and which are functionally related to transcriptional activity (Fig. 5b, c). A ~5.5-fold enrichment of CENP-A^TAP^ was found at these sites (Fig. 5d). CENP-A assembled into chromatin when expressed at endogenous levels was also found to be enriched 3-fold at DNase I hypersensitive sites (Fig. 5e) and promoters (Fig. S3f). Conversely, both CENP-A^TAP^ and CENP-A^LAP^ were not enriched at H3k27me3 peak sites that are tightly associated with inactive gene promoters (Fig. S3c, e).

### Ectopic CENP-A is removed contemporaneously with replication fork progression, while centromeric CENP-A is retained

We next tested whether removal by G2 of CENP-A assembled into nucleosomes at unique sites on the chromosome arms is mediated by the direct action of the DNA replication machinery. CENP-A^TAP^ was affinity purified from mid S phase cells and CENP-A-bound DNAs were sequenced (Fig. 6a; Fig. S4a). In parallel, we pulse-labeled newly synthesized DNA in our synchronized cells by addition of bromodeoxyuridine (BrdU) for 1 hour at early (S0-S1), mid (S3-S4), and late S phase (S6-S7) (Fig. 6a; Fig. S4a). Genomic DNA from each time point was sonicated (Fig. S4b) and immunoprecipitated with a BrdU antibody (Fig. 6a). Eluted DNA was then sequenced and mapped to the genome [an approach known as Repli-seq ^56^], yielding regions of early, mid, and late replicating chromatin (an example from a region of chromosome 20 arm is shown in Fig. 6b). The early replication timing was validated (using qPCR - Fig. S4c) for two genes (MRGPRE and MMP15) previously reported to be early replicating [ref ^57^ and ENCODE Repli-seq]. Similarly, a gene and a centromeric region (HBE1 and Sat2) previously reported to be late replicating [ref ^58^ and ENCODE Repli-seq] were confirmed in our cells to be replicated late (Fig. S4d).

**Figure 6.**
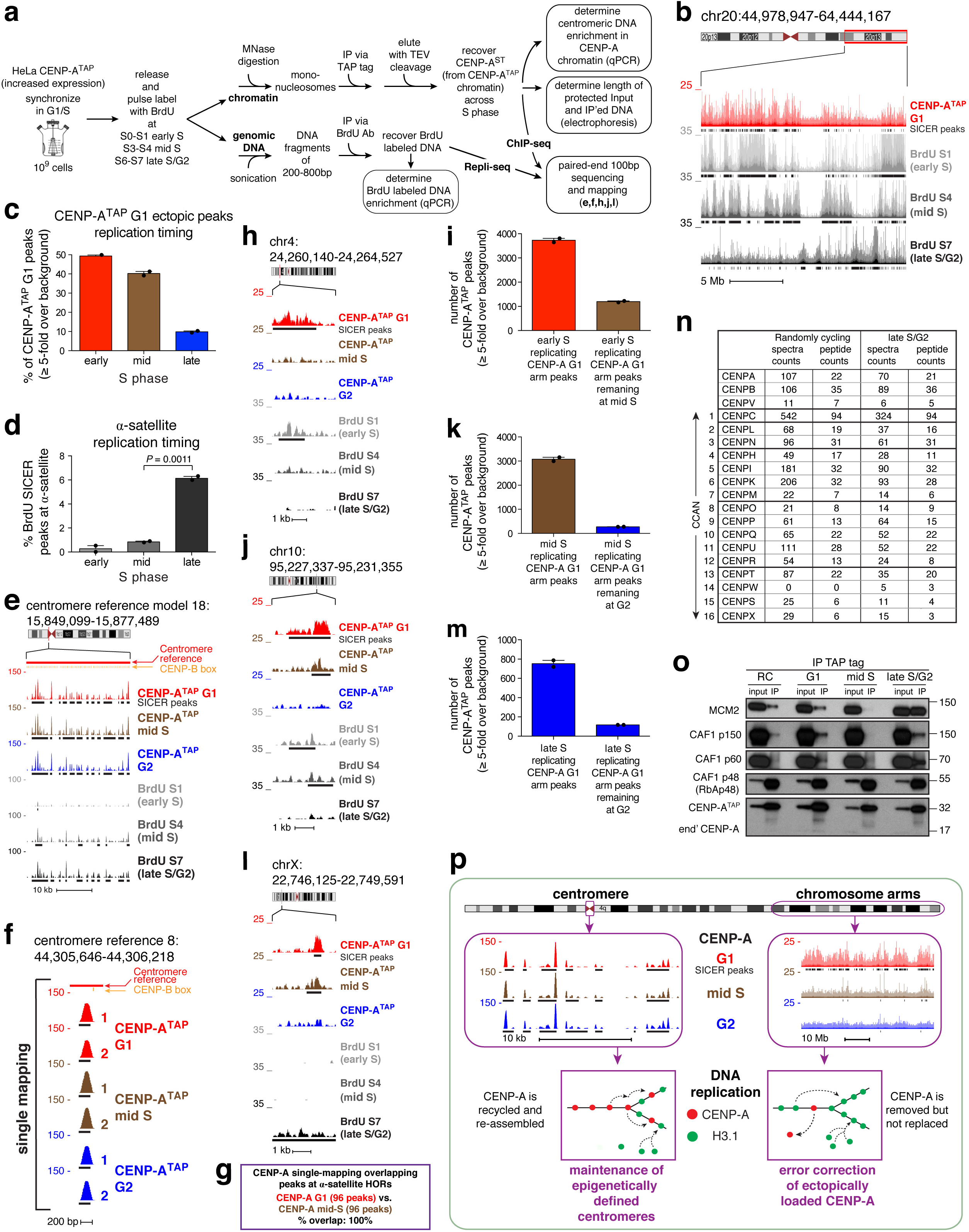
Ectopic CENP-A is removed contemporaneously with replication fork progression, while centromeric CENP-A is retained. (**a**) Schematic representation of CENP-A ChIP-seq combined with Repli-seq experiment across S phase. (**b**) Raw mapping data of CENP-A^TAP^ ChIP-seq at G1 (red) and BrdU repli-seq (grey scale) at early S (S1), mid S (S4) and late S/G2 (S7) at the q-arm of chromosome 20. SICER peaks, black lines are drawn underneath the raw mapping data. Scale bar, 5 Mb. (**c**) The percentage of ectopic G1 CENP-A^TAP^ ≥5-fold binding sites that are found in early, mid, or late S replicating regions. N=2 for from two independent replicates. Error bars, s.e.m. (**d**) Percentage of BrdU SICER peaks at α-satellite DNA found within early, mid or late S replicating regions. N=2 for from two independent replicates. Error bars, s.e.m. *P* value of 0.0011 determined using two-tailed *t*-test. (**e**) CENP-A ChIP-seq raw mapping data at a part of cen18 at G1, mid S phase and G2, and BrdU repli-seq at early S (S1), mid S (S4) and late S/G2 (S7). SICER peaks are denoted as black lines underneath the raw mapping data. Centromere reference location, red. CENP-B boxes, orange. Scale bar, 10kb. (**f**) High-resolution view of CENP-A mapping during DNA replication (mid-S) to a single copy site (marked by a purple bar in Fig. 2a) in the centromere reference model of chromosome 8. Data from Figure 2c for CENP-A^TAP^ G1 and G2 is included for comparison. Scale bar, 200bp. (**g**) Complete overlap between CENP-A G1 and mid-S single mapping binding sites (after filtering out multi-mapping reads) is found at α-satellite HORs sequences. (**h, j, l**) Raw mapping data (colored) and SICER peaks (black lines, underneath) of CENP-A^TAP^ ChIP-seq at G1, mid S phase and G2 and BrdU labeled repli-seq samples (grey scale) showing regions going through replication at early S (S1), mid S (S4) and late S/G2 phase (S7) within the p-arm of chromosome 4 (**h**), the q-arm of chromosome 10 (**j**), and the p-arm of chromosome X. Early replicating CENP-A^TAP^ G1 peaks (**h, j**) are removed by mid S phase. Mid-S replicating CENP-A^TAP^ G1 (**j**) peaks are removed by G2. Late replicating CENP-A^TAP^ G1 peaks (**l**) are removed by G2. Scale bars, 1kb. (**i**) Early replicating CENP-A^TAP^ G1 peaks (≥5-fold over background - shown in (**h**) for chromosome 4) were analyzed for their retention at mid S phase. N=2 for from two independent replicates. Error bars, s.e.m. About 70% of early replicating CENP-A^TAP^ G1 peaks are removed by mid S phase. (**k**) Mid-S replicating CENP-A^TAP^ G1 peaks (≥5-fold over background - shown in (**j**) for chromosome 10) were analyzed for their retention at G2. N=2 for from two independent replicates. Error bars, s.e.m. 90% of mid replicating CENP-A peaks are removed by G2. (**m**) Late replicating CENP-A^TAP^ G1 peaks (≥5-fold over background - shown in (**l**) for chromosome X) were analyzed for their retention at G2. N=2 for from two independent replicates. Error bars, s.e.m. 85% of late replicating CENP-A^TAP^ G1 peaks are removed by G2. (**n**) CENP-A^TAP^ was immunoprecipitated from the chromatin fraction of randomly cycling cells or late S/G2 synchronized cells followed by mass spectrometry to identify the co-precipitated partners. All the CCAN network components were co-precipitated with CENP-A at late S/G2. (**o**) CENP-A^TAP^ was immunoprecipitated from micrococcal nuclease resistant chromatin isolated at different cell cycle phases and the immunoprecipitates were examined by immunoblotting for CAF1 complex subunits and MCM2. (**p**) Model for maintaining centromeric CENP-A while removing it from non-centromeric sites on the chromosome arms during DNA replication to ensure maintenance of centromere identity across the cell cycle.

CENP-A immunoprecipitation from micrococcal nuclease digested chromatin isolated from early and mid S phase cells resulted in levels of α-satellite DNA enrichment (Fig. S4e) similar to those achieved at G1 phase (Fig. 1b). Furthermore, nucleosomal CENP-A chromatin produced by micrococcal nuclease digestion protected 133 bp of DNA at early and mid S phase (Fig. S4f) just as it did in G1 and G2 [Fig. 1c; see also ref ^8^], with no evidence for a structural change from hemisomes to nucleosomes and back to hemisomes during S phase as previously claimed ^59^. Mapping of CENP-A binding sites within the chromosome arms, combined with Repli-seq analysis, revealed that 91% of ectopic G1 CENP-A binding sites were found in early- or mid-S replicating regions (Fig. 6b, c). While centromeres of chromosomes 1, 3, 10, 17, 18 and X ^60^ and bulk α-satellite DNAs or a consensus pool of alphoid DNA sequences have been reported to replicate at mid ^61, 62^ or mid-to-late ^63^ S phase, in our cells α-satellite containing DNAs in all 23 centromeres were found almost exclusively to be late replicating (Fig. 6d).

Remarkably, throughout S phase, centromere bound CENP-A found in G1 was completely retained across each reference centromere with the same sequence binding preferences (shown for centromere 18 in Fig. 6e and Fig. S4g). Retention of CENP-A binding during DNA replication was observed also at the unique sequence binding sites within HORs (Fig. 6f). Indeed, all 96 CENP-A^TAP^ G1 peaks at single copy variants within α-satellite HORs remained bound by CENP-A at mid S phase (Fig. 6g). In contrast, early replicating ectopic CENP-A binding sites were nearly quantitatively removed during or quickly after their replication and were no longer visible at mid-S phase (Fig. 6h, i). Similarly, ectopic CENP-A binding sites that were in mid-S replicating regions remained at mid-S but were removed quickly after that and were absent by late S/G2 (Fig. 6j, k). Ten percent of ectopic CENP-A G1 peaks were in late-S replicating regions (Fig. 6c). Here again, almost all (85%) of these were removed by G2 (Fig. 6l, m), while late replicating centromeric CENP-A peaks were retained (Fig 6d-g). These results demonstrate that ectopic, but not centromeric, CENP-A binding sites are removed as DNA replication progresses. Moreover, that contemporaneously late replicating centromeric bound CENP-As, including the unique binding sites within the 23 centromeres, were retained following DNA replication while ectopic sites were removed eliminates the possibility that retention could be a consequence of a general alteration in late S phase in the activity of one or more DNA replication components that could potentially act to facilitate CENP-A reloading.

### Centromeric CENP-A is continuously bound by the CCAN complex during centromeric DNA replication

To comprehensively determine the components which associate with CENP-A during replication in late S, we used mass spectrometry following affinity purification of CENP-A nucleosomes (Fig. S4h, left panel). A structural link that normally bridges multiple centromeric CENP-A nucleosomes and nucleates full kinetochore assembly before mitotic entry ^64–70^ is the 16-subunit <u>c</u>onstitutive <u>c</u>entromere <u>a</u>ssociated <u>n</u>etwork (CCAN). This complex is anchored to CENP-A primarily through CENP-C ^68, 71–75^ and sustained by CENP-B binding to α-satellites ^76^.

Remarkably, mass spectrometry identified that all 16 CCAN components ^23, 25^ remained associated with mono-nucleosomal CENP-A chromatin affinity purified from late S/G2 (Fig.6n; Fig. S4h). Further, association of CENP-A with MCM2 (and other components of the MCM2–7 replicative helicase complex) and CAF1p150 was enhanced at late S phase (compared with its association in randomly cycling cells) (Fig. S4i). Stable association with CENP-A was also seen for HJURP, multiple chromatin remodeling factors and nuclear chaperones (Fig. S4k), histones (Fig. S4l), centromere and kinetochore components (Fig. S4m), and other DNA replication proteins (Fig. S4j).

The continuing interaction during DNA replication of CCAN proteins with CENP-A and which is maintained even on mono-nucleosomes provides strong experimental support that the CCAN network tethers CCAN-bound centromeric CENP-A at or near the centromeric DNA replication forks, thereby enabling its efficient reincorporation after replication fork passage. To test this further, the composition of CENP-A-containing nucleosomal complexes from G1 to late S/G2 was determined following affinity purification (via the TAP tag) of chromatin-bound CENP-A^TAP^ from predominantly mononucleosome pool (Fig. S4h, right panel).

We initially focused on the <u>C</u>hromatin <u>A</u>ssembly <u>F</u>actor 1 (CAF1) complex, which is required for *de novo* chromatin assembly following DNA replication ^77, 78^. Its p48 subunit (also known as CAF1 subunit c, RbAp48, or RBBP4) 1) binds histone H4 ^79^ and 2) has been reported as a binding partner in a CENP-A pre-nucleosomal complex with HJURP and nucleophosmin (NPM1) ^34^. In this latter complex it has been proposed to promote H4K5 and K12 acetylation prior to CENP-A loading ^80^ and maintain the deacetylated state of histones in the central core of centromeres after deposition ^81^. Immunoblotting revealed that CAF1 p48 co-immunopurified with CENP-A from G1 through late S/G2 (Fig. 6o), consistent with a role for it in binding H4 and perhaps maintaining a deacetylated state.

In striking contrast, the two other CAF1 subunits, CAF1 p150 and CAF1 p60, that are essential for *de novo* chromatin assembly *in vitro* ^82^, remained much more strongly associated with CENP-A nucleosomes in late S/G2 than in mid-S (Fig. 6o). Additionally, MCM2, a core subunit of the DNA replicative helicase MCM2–7 complex that has an important role in recycling of old histones as the replication fork advances ^83^, was robustly co-purified with CENP-A only in late S phase derived chromatin, with no association detected in mid-S (Fig. 6o), when ectopic CENP-A peaks replicate. Thus, there is stable association of CENP-A with MCM2 and the CAF1 subunits necessary for chromatin reassembly after replication only in late S phase, the time when all centromeric, but only a small minority of ectopically loaded, CENP-A is replicated.

## Discussion

Using reference models for 23 human centromeres, we have identified that during DNA replication CENP-A nucleosomes initially assembled onto centromeric α-satellite repeats are reassembled onto the same spectrum of α-satellite repeat sequences of each daughter centromere as was bound prior to DNA replication. Additionally, genome-wide mapping of sites of CENP-A assembly identified that when CENP-A is expressed at endogenous levels, the selectivity of the histone chaperone HJURP’s loading in early G1 of new CENP-A at or near existing sites of centromeric CENP-A-containing chromatin is insufficient to prevent its loading onto >11,000 sites along the chromosome arms (Fig. 3d). We also show that the number of ectopic sites increases as CENP-A expression levels increase, as has been reported in multiple human cancers ^39, 63, 64^. These sites of ectopic CENP-A are replicated in early and mid-S (Fig. 6b, c) and are nearly quantitatively removed as DNA replication progresses (Fig. 6h-m).

Taken together, our evidence identifies that DNA replication functions not only to duplicate centromeric DNA but also as an error correction mechanism to maintain epigenetically-defined centromere position and identity by coupling centromeric CENP-A retention with its removal from assembly sites on the chromosome arms (Fig. 6p). Indeed, our data reveal that CENP-A loaded onto unique sites (after filtering out multi-mapping reads in the α-satellite HORs in the HuRef genome) within the 23 reference centromeres, is precisely maintained at these sites during and after DNA replication, offering direct support that at least for each of these unique sites the replication machine re-loads CENP-A back onto the exact same centromeric DNA site (Fig. 2, 6f, g). Accompanying this is retention of centromeric, α-satellite DNA-bound CENP-A before and after DNA replication at indistinguishable sequences throughout reference models of all 23 human centromeres (Fig. S2). DNA replication produces a very different situation for CENP-A initially assembled into nucleosomes on the chromosome arms. Sites of this ectopically loaded CENP-A are nearly quantitatively stripped during DNA replication (Figs. 3, 4, 6h-m), providing strong evidence that DNA replication acts not only to duplicate both centromeric and non-centromeric DNA sequences, but also to reinforce epigenetically defined centromere position and identity, while precluding acquisition of CENP-A-dependent centromere function at non-centromeric sites (Fig. 6p).

Without such correction, ectopically loaded sites would be maintained and potentially reinforced cell cycle after cell cycle, ultimately recruiting CENP-C which in turn can nucleate assembly of the CCAN complex ^23–26^. Increasing levels of arm-associated CENP-A/CCAN would present a major problem for faithful assembly and function of a single centromere/kinetochore per chromosome, both by acquisition of partial centromere function and by competition with the authentic centromeres for the pool of available CCAN components. Indeed, high levels of CENP-A overexpression 1) leads to recruitment of detectable levels of 3 of 16 CCAN components (CENP-C, CENP-N and Mis18) assembled onto the arms ^36, 37, 84^, 2) ongoing chromosome segregation errors ^38^, and 3) has been observed in several cancers where it has been associated with increased invasiveness and poor prognosis ^39, 85, 86^.

As to the mechanism for retention during DNA replication of centromeric but not ectopically loaded CENP-A, an attractive model strongly supported by our evidence is that the local reassembly of CENP-A within centromeric domains is mediated by the continuing centromeric CENP-A association with CCAN complexes, which we show to be maintained on individual CENP-A nucleosomes in late S when centromeres are replicated (Fig. 6n). The continued presence of the assembled CCAN network directly bound to CENP-A during centromere DNA replication offers a plausible explanation for centromeric CENP-A retention (together with the MCM2 replicative helicase and the major CAF1 subunits required for nucleosome reassembly post-replication ^77, 83^). In such a model, centromere identity is preserved by an assembled CCAN network which serves during DNA replication to tether disassembled CENP-A/H4 dimers or tetramers near the sites of centromere replication, thereby enabling their local reassembly onto each of the daughter centromeres and the corresponding epigenetic inheritance of centromere identity.

## Supplementals figure legends

**Supplementary Figure S1.**
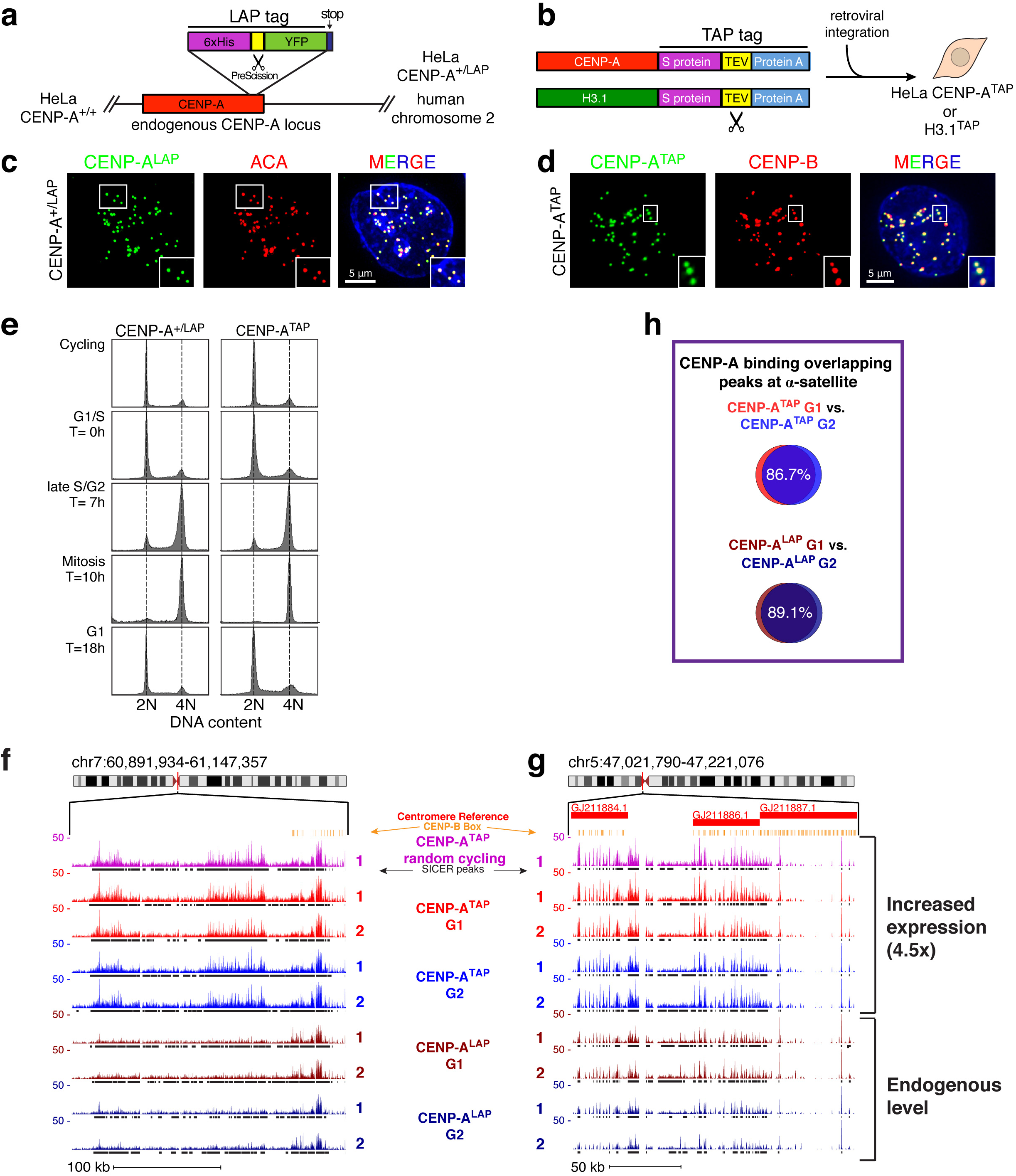
Identification of peaks enriched for CENP-A binding. **(a)** Scheme showing experimental design for tagging an endogenous CENP-A locus to produce CENP-A^+/LAP^ HeLa cells. These cells were then adapted to suspension growth. (**b**) Scheme showing the experimental design for obtaining increased levels of CENP-A^TAP^ expression. CENP-A^TAP^ is expressed in these cells at 4.5-fold the level of CENP-A in the parental HeLa cells ^8^. (c, d) Localization of endogenously tagged CENP-A^LAP^ (c) and CENP-A^TAP^ (**d**) determined with indirect immunofluorescence using anti-GFP antibody (**c**) or rabbit-IgG (**d**). Scale bar, 5 μm.(**e**) FACS analysis of DNA content showing the synchronization efficiency of CENP-A^+/LAP^ and CENP-A^TAP^ HeLa cell lines. (**f**, **g**) Examples of centromeric regions of chromosome 7 (**f**) and 5 (**g**) showing increased occupancy of overexpressed CENP-A^TAP^ (compare CENP-A^TAP^ with CENP-A^LAP^). (**h**) Overlap between G1 and G2 CENP-A binding peaks at α-satellite sequences.

**Supplementary Figure S2.**
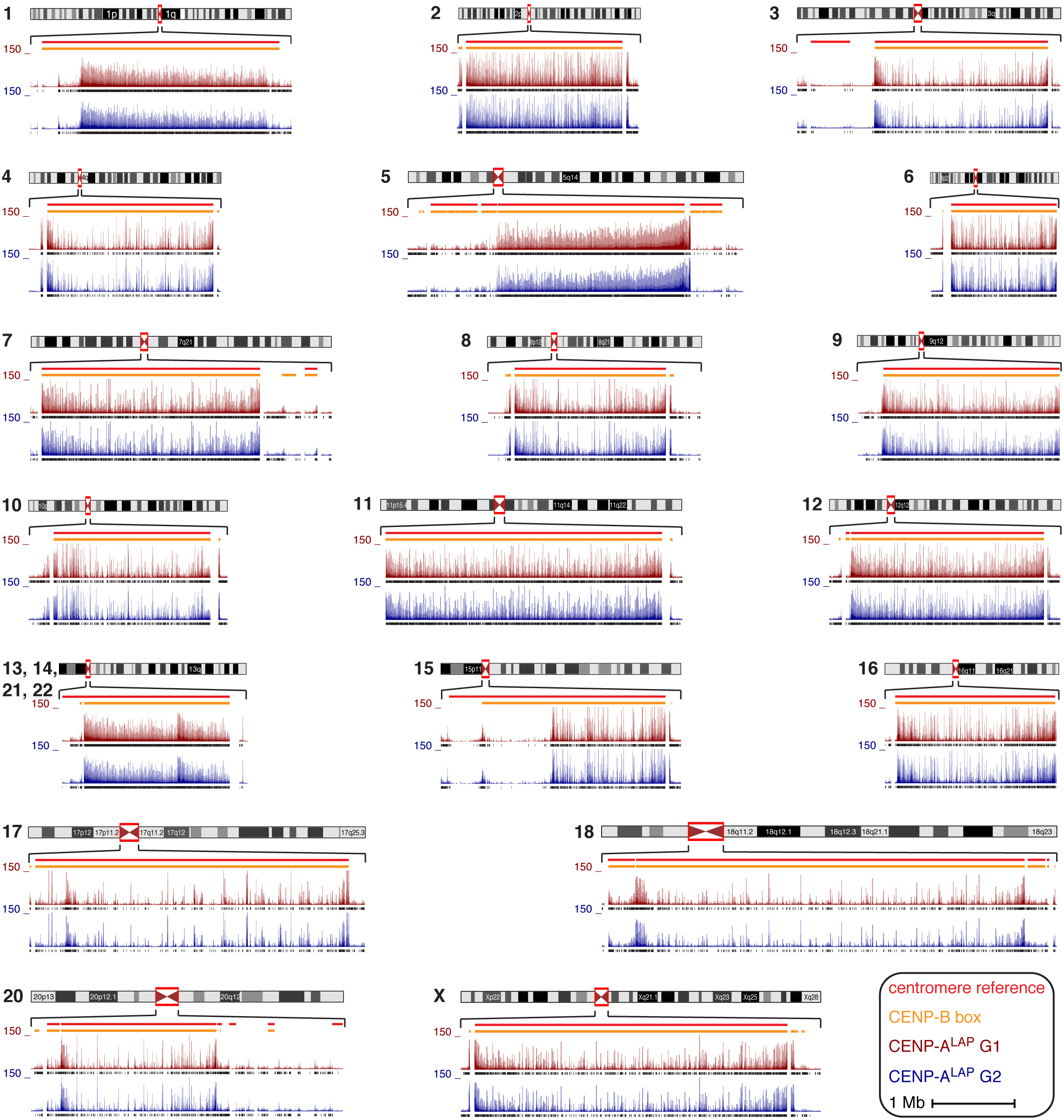
CENP-A ChIP-seq identifies CENP-A binding at reference centromeres of 23 human chromosomes. CENP-A^LAP^ bound DNAs at G1 and G2 were sequenced, with 2 replicates per condition, and mapped to the centromeric reference models in the hg38 assembly ^46^. Shown are the raw mapping data (colored) for every human centromere (except for the centromere of chromosome 19 that shares almost all of its α-satellites arrays with α-satellites arrays of chromosomes 1 and 5) and CENP-A binding called as SICER peaks (black lines, underneath) for one replicate for each time point. Centromere reference location, red. CENP-B box, orange.

**Supplementary Figure S3.**
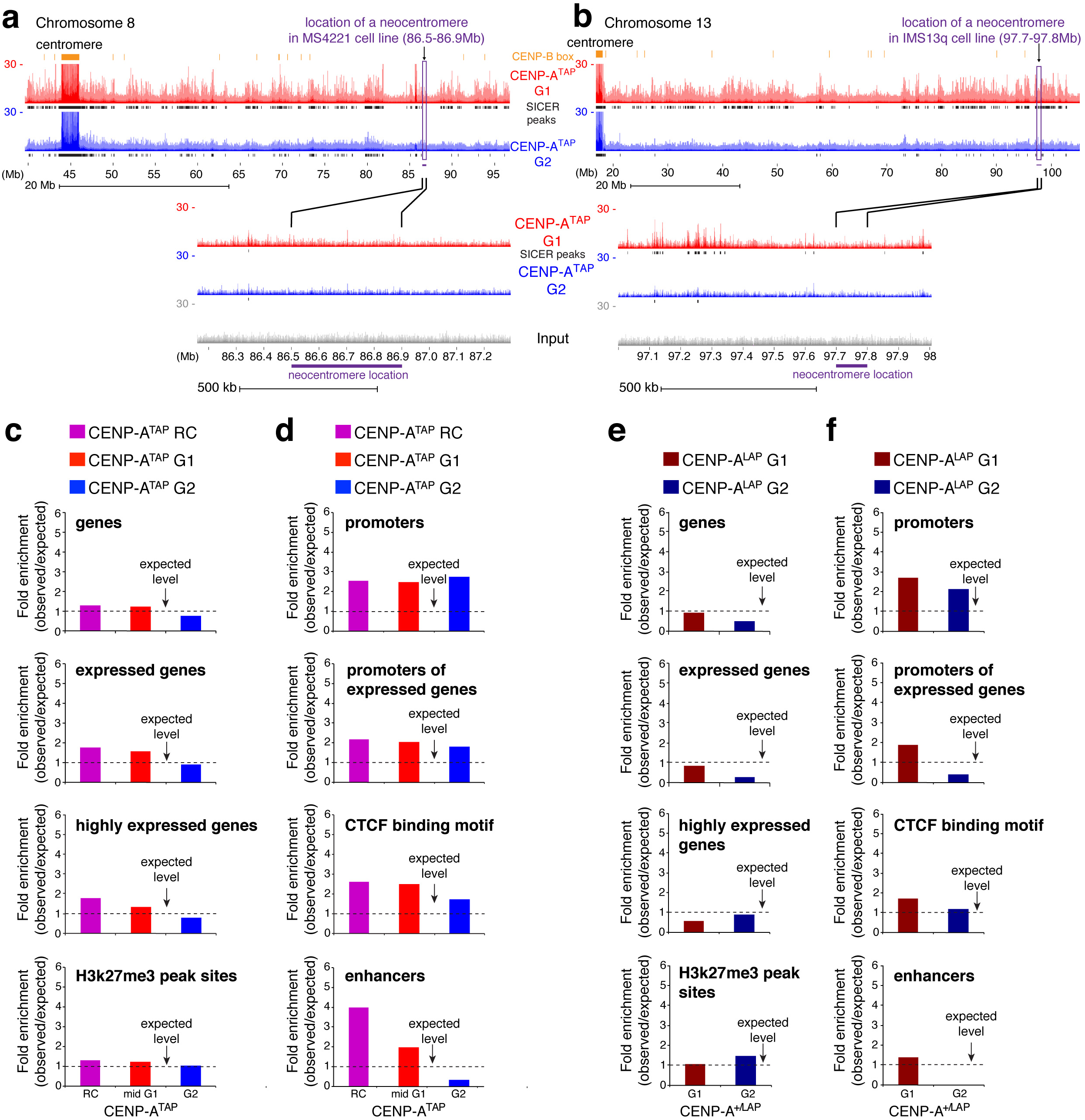
Ectopic deposition of CENP-A into open and active chromatin at G1 does not function as a seeding hotspot for neocentromere formation. (**a, b**) Read mapping data of CENP-A^TAP^ ChIP-sequencing at G1 (red) and G2 (blue), at the chromosomal location of 2 known patient derived neocentromeres ^28^ found in chromosome 8 (**a**) and chromosome 13 (**b**). **(c-f**) Fold enrichment of CENP-A^TAP^ chromatin in randomly cycling cells or at G1 or G2 (**c, d**) and CENP-A^LAP^ chromatin (**e, f**) at G1 or G2 at different genomic locations. SICER peaks ≥5-fold supported between two replicates were analyzed for their enrichment level at different genomic locations, compared to the level of enrichment at these sites by chance.

**Supplementary Figure S4.**
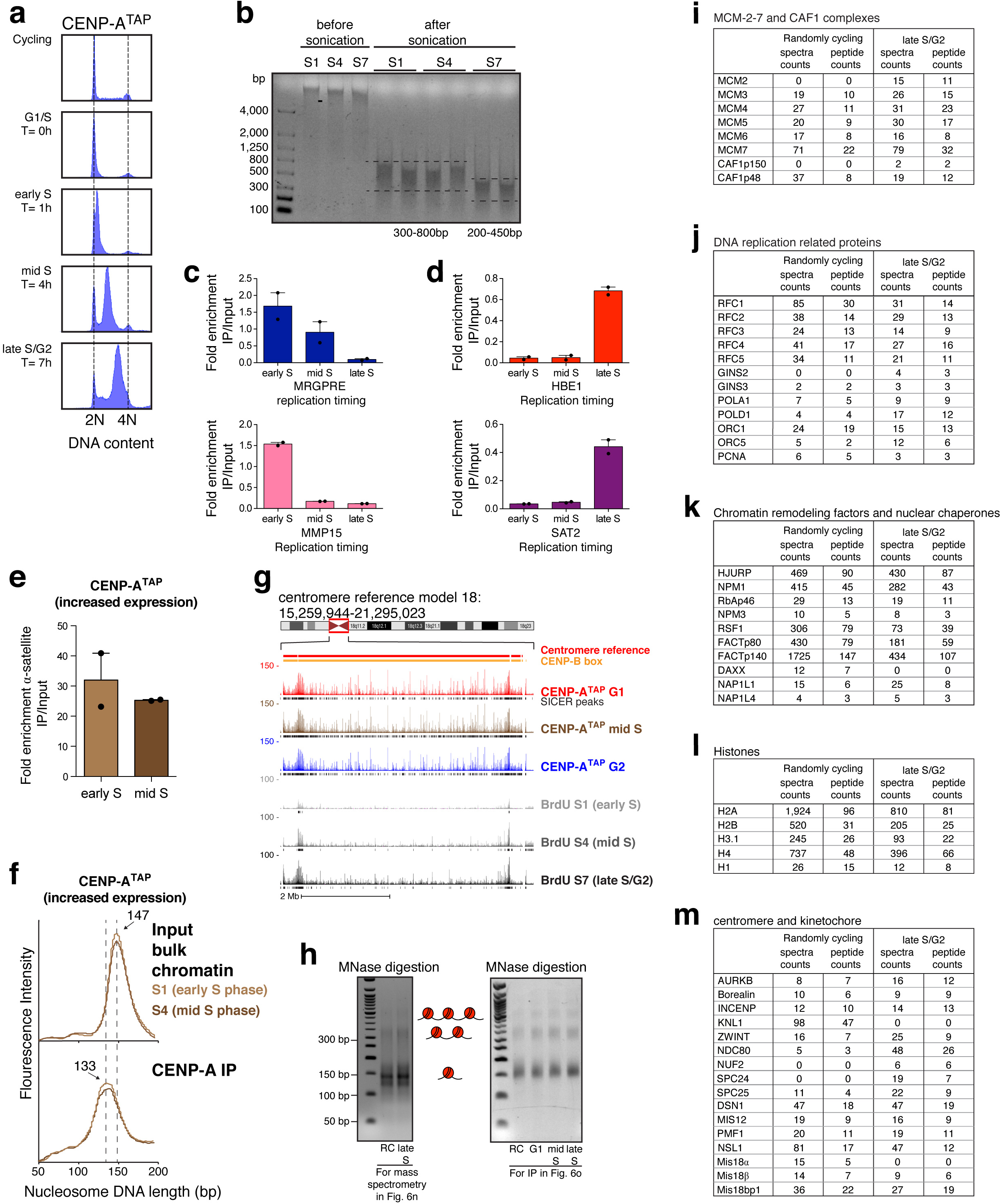
Centromeres are late replicating with CENP-A remaining tethered locally by continued binding to the CCAN complex. (**a**) FACS analysis of DNA content showing the synchronization efficiency of CENP-A^TAP^ HeLa cell line across S phase. (b) Genomic DNA of cells labeled for 1 hour with BrdU was sonicated prior to the BrdU immunoprecipitation and fragments of 200–800bp were obtained. (**c**) Quantitative real-time PCR for MRGPRE and MMP15 validates their early replication timing as previously reported [ref ^57^ and ENCODE Repli-seq]. N=2, from two independent replicates. Error bars, s.e.m. (**d**) Quantitative real-time PCR for HBE1 and Sat2 was used to validate their late replication timing, as previously reported [ref ^58^ and ENCODE Repli-seq]. N=2, from two independent replicates. Error bars, s.e.m. (**e**) Quantitative real-time PCR for α-satellite DNA. N=2, from two independent replicates. Error bars, s.e.m. (**f**) MNase digestion profile showing the nucleosomal DNA length distributions of bulk input mono-nucleosomes (upper panel) and purified CENP-A^TAP^ following native ChIP at early S and mid S phase. (**g**) CENP-A ChIP-seq raw mapping data spanning the whole of cen18 at G1, mid S phase and G2, and BrdU repli-seq at early S (S1), mid S (S4) and late S/G2 (S7). SICER peaks are denoted as black lines underneath the raw mapping data. Centromere reference location, red. CENP-B boxes, orange. Scale bar, 2Mb. (**h**) Ethidium Bromide stained DNA agarose gel showing MNase digestion profile of bulk chromatin used for mass spectrometry identification of proteins associating with CENP-A^TAP^ chromatin (left panel) and for CENP-A^TAP^ co-immunoprecipitation experiment (right panel). (**i-m**) CENP-A^TAP^ immunopurification followed by mass spectrometry identifies association with CENP-A chromatin of DNA replication related proteins (**i, j**), chromatin remodeling factors and nuclear chaperones (**k**), histones (**l**) and centromere and kinetochore proteins (**m**).

## Materials and Methods

### Cell lines

Adherent HeLa cells stably expressing CENP-A^TAP^ or H3.1^TAP^ by retrovirus infection ^23^ or endogenously tagged CENP-A^+/LAP^ by infection of a rAAV harboring a LAP targeting construct containing homology arms for CENP-A ^43^ were adapted to suspension growth by selecting surviving cells and were maintained in DMEM medium (Gibco) containing 10% fetal bovine serum (Omega Scientific), 100U/ml penicillin, 100U/ml streptomycin and 2mM l-glutamine at 37°C in a 5% CO_2_ atmosphere with 21% oxygen. Cells were maintained and split every 4–5 days according to ATCC recommendations.

### Cell synchronization

Cells were synchronized as previously described ^8^. Briefly, suspension HeLa cells were treated with 2 mM thymidine in complete medium for 19 h, pelleted and washed twice in PBS, and released in complete medium containing 24 μM deoxycytidine for 9 h followed by addition of thymidine to a final concentration of 2 mM for 16 h, after which cells were released again into complete medium containing 24 μM deoxycytidine. For G2, cells were harvested 7 hours after release from the second thymidine block. For G1, thymidine was added for a third time, 7 hours after the release and cells were harvested 11 hours after that (a total of 18 hours after the release from the second thymidine block).

### Chromatin extraction

Chromatin was extracted Nuclei from 1×10^9^ nuclei of HeLa cells as previously described ^8^. Nuclei from 1×10^9^ HeLa cells were prepared by pelleting and resuspending cells in buffer containing 3.75 mM Tris at pH 7.5, 20 mM KCl, 0.5 mM EDTA, 0.5 mM DTT, 0.05 mM spermidine, 0.125 mM spermine, 1 mM PMSF and 0.1% digitotin. Cells were homogenized with 10 strokes and nuclei were pelleted at 300*g*. Nuclei were then washed once in wash buffer (20 mM HEPES at pH 7.7, 20 mM KCl, 0.5 mM EDTA, 0.5 mM DTT and 0.5 mM PMSF), followed by wash buffer containing 150 mM NaCl. Nuclei were resuspended in wash buffer supplemented with 150 mM NaCl and 3 mM CaCl_2_. Chromatin was digested at room temperature using 140 units ml–1 of micrococcal nuclease (Roche, 10107921001) for 35 minutes to produce a pool of mono-nucleosomes. Following micrococcal nuclease treatment, extracts were supplemented with 5 mM EGTA and 0.05% NP40 and centrifuged at 10,000g for 15 min at 4 °C. The supernatant was then used as the starting material for all immunopurifications.

### Affinity purification

TAP-or LAP -tagged chromatin were purified in two steps. In the first step, native TAP-tagged chromatin was immunoprecipitated by incubating the bulk soluble mono-nucleosome pool with rabbit IgG (Sigma-Aldrich) coupled to Dynabeads M-270 Epoxy (Thermo Fisher Scientific, 14301). Alternatively, CENP-A^LAP^ chromatin was immunoprecipitated using mouse anti-GFP antibody (clones 19C8 and 19F7, Monoclonal Antibody Core Facility at Memorial Sloan-Kettering Cancer Center, New York) ^87^ coupled to Dynabeads M-270 Epoxy. Chromatin extracts were incubated with antibody bound beads for 16 h at 4 °C. Bound complexes were washed once in buffer A (20 mM HEPES at pH 7.7, 20 mM KCl, 0.4 mM EDTA and 0.4 mM DTT), once in buffer A with 300 mM KCl and finally twice in buffer A with 300 mM KCl, 1 mM DTT and 0.1% Tween 20. In the second step, TAP-chromatin complexes were incubated 16 h in final wash buffer with 50μl recombinant TEV protease, resulting in cleavage of the TAP tag and elution of the chromatin complexes from the beads. Alternatively, CENP-A^LAP^ chromatin was eluted from the beads by cleaving the LAP tag using PreScission protease (4 h, 4°C).

### DNA extraction

Following elution of the chromatin from the beads, Proteinase K (100 μg/ml) was added and samples were incubated for 2 h at 55°C. DNA was purified from proteinase K treated samples using a DNA purification kit following the manufacturer instructions (Promega, Madison, USA) and was subsequently analyzed either by running a 2% low melting agarose (APEX) gel or by an Agilent 2100 Bioanalyzer by using the DNA 1000 kit. The Bioanalyzer determines the quantity of DNA on the basis of fluorescence intensity.

### Quantitative real-time PCR (qPCR)

Quantitative real-time PCR (qPCR) was performed using SYBR Green mix (Bio Rad) with CFX384 Bio Rad Real Time System. Primers sequences used in this study: MRGPRE: (forward) 5’-CTGCGCGGATCTCATCTTCC-3’ and (reverse) 5’-GGCCCACGATGTAGCAGAA-3’. MMP15: (forward) 5’-GTGCTCGACGAAGAGACCAAG-3’ and (reverse) 5’-TTTCACTCGTACCCCGAACTG-3’. HBE1: (forward) 5’-ATGGTGCATTTTACTGCTGAGG-3’ and (reverse) 5’-GGGAGACGACAGGTTTCCAAA-3’. Sat2: (forward) 5’-TCGCATAGAATCGAATGGAA-3’ and (reverse) 5’-GCATTCGAGTCCGTGGA-3’ ^16^. α-satellite DNA (from chromosomes 1, 3, 5, 10, 12 and 16): (forward) 5’-CTAGACAGAAGAATTCTCAG-3’ and (reverse) 5’-CTGAAATCTCCACTTGC-3’ ^55^. Melting curve analysis was used to confirm primer specificity. To ensure linearity of the standard curve, reaction efficiencies over the appropriate dynamic range were calculated. Using the dCt method, we calculated fold-enrichment of α-satellite DNA after immunopurification of CENP-A^TAP^ chromatin, compared to its level in the bulk input chromatin. For Repli-seq experiment, we used the dCt method, to calculate fold-enrichment of replicated DNA after immunopurification of BrdU-labeled DNA compared to its level in the bulk input DNA. Reported values are the means of two independent biological replicates with technical duplicates that were averaged for each experiment. Error bars represent SE of the mean.

### Immunoblotting

For immunoblot analysis, protein samples were separated by SDS–PAGE, transferred onto PVDF membranes (Millipore) and then probed with the following antibodies: rabbit anti-CENP-A (Cell Signaling, 2186s, 1:1,000), rabbit anti-CENP-B (Millipore, 07–735, 1:200), mouse anti-α-tubulin (Abcam, DM1A, 1:5000), rabbit anti-CAF1p150 (Santa Cruz, sc-10772, 1:500), rabbit anti-CAF1p60 (Bethyl Laboratories, A301–085A, 1:1,000), rabbit anti-CAF1p48 (Bethyl Laboratories, A301–206A, 1:1,000), rabbit anti-MCM2 (Abcam, Ab4461, 1:1,000). Following incubation with HRP-labelled antibody (GE Healthcare, NA931V or NA934V), HRP was detected using enhanced chemiluminescence (ECL) substrate (Thermo Scientific, 34080 or 34096).

### Immunofluorescence

1×10^6^ suspension cells were centrifuged and resuspended with PBS. 10^5^ cells were immobilized on glass slide by cytospin centrifugation for 3 min, 800rpm. Cells were then fixed using ice-cold methanol at −20°C for 10 min, followed by washing with cold PBS and then incubated in Triton Block (0.2 M glycine, 2.5% FBS, 0.1% Triton X-100, PBS) for one hour. The following primary antibodies were used: mouse anti-GFP (Roche, 11814460001, 1:500), rabbit anti-CENP-B (Abcam 25734, 1:1,000), human anti-centromere antibodies (ACA, Antibodies Inc, 15–234–0001, 1:500). The following secondary antibodies (Jackson Laboratories) were used for 45 minutes: donkey anti-human TR (1:300), anti-mouse FITC (1:250). TAP fusion proteins were visualized by incubation with FITC-rabbit IgG (Jackson Laboratories, 1:200). Cells were then washed with 0.1% Triton X-100 in PBS, counterstained with DAPI and mounted with mounting medium (Molecular Probes, P36934). Immunofluorescent images were acquired on a Deltavision Core system at x60–100 magnification. 0.2 μm *Z*-stack deconvolved projections were generated using the softWoRx program.

### Flow cytometry

Flow cytometry was used to determine the DNA content of the cells as. 1×10^6^ cells were harvested, washed in PBS and fixed in 70% ethanol. Cells were then washed and DNA was stained by incubating cells for 30 min with 1% FBS, 10 μg ml^-1^ propidium iodide and 0.25 mg ml^-1^ RNase A in PBS followed by FACS analysis for DNA content using a BD LSR II Flow Cytometer (BD Biosciences).

### ChIP-Seq Library Generation and Sequencing

ChIP libraries were prepared following Illumina protocols with minor modifications (Illumina, San Diego, CA). To reduce biases induced by PCR amplification of a repetitive region, libraries were prepared from 80–100 ng of input or ChIP DNA. The DNA was end-repaired and A-tailed and Illumina Truseq adaptors were ligated. Libraries were run on a 2% agarose gel. Since the chromatin was digested to mononucleosomes, following adaptors ligation the libraries size was 250–280 bp. The libraries were size selected for 200–375 bp. The libraries were then PCR-amplified using only 5–6 PCR cycles since the starting DNA amount was high. Resulting libraries were sequenced using 100 bp, paired-end sequencing on a HiSeq 2000 instrument per manufacturer’s instructions with some modifications (Illumina, San Diego, CA). Sequence reads are summarized in Table S1.

### Initial sequence processing and alignment

Illumina paired-end reads were merged to determine CENP-A or H3 containing target fragments of varying length using PEAR software ^88^, with standard parameters (p-value: 0.01, min-overlap: 10 bases, min-assembly length: 50bp). Merged paired reads were mapped (Bwa-Mem, standard parameters ^89, 90^) to the human genome 38 (hg38) assembly (including alternative assemblies), which contain human α-satellite sequence models in each centromeric region (^45^; BioProject: PRJNA193213; ^46^). Reads were determined to contain α-satellite if they overlapped sites (BEDTools: intersect ^91^) in the genome previously annotated as α-satellite (UCSC table browser ^92^ was used to obtain a bed file of all sites annotated as ALR/α-Satellite). Additionally, merged sequences were defined as containing α-satellite if they contained an exact match to at least two 18-mers specific to a previously published WGS read database of α-satellite, representing 2.6% of sequences from the HuRef genome ^21, 48^. Comparisons between the Bwa mapping and 18-mer exact matching based strategies were highly concordant. Total α-satellite DNA content in human genome 38 assembly was estimated by using the UCSC RepeatMasker Annotation ^92, 93^. Summary of reads obtained is shown in Table S1.

### ChIP-seq peak calling

Enrichment peaks for ChIP experiments were determined using SICER algorithm (v1.0.3) ^50^ using relevant input reads as background, with stringent parameters previously optimized for human CENP-A ^36^: threshold for redundancy allowed for chip reads: 1, threshold for redundancy allowed for control reads: 1, window size: 200 bps, fragment size: 150 bps, shift in window length is 150, effective genome size as a fraction of the reference genome of hg38: 0.74, gap size: 400 bps, e-value for identification of candidate islands that exhibit clustering: 1000, and false discovery rate controlling significance: 0.00001. In parallel, MACs peak calling was performed (macs14) ^49^, and wiggle tracks were created to represent read depth of each dataset independently. Finally, we performed a final, rigorous evaluation of ectopic CENP-A peaks, or peaks predicted outside of centromeric regions, using k-mer enrichment (previously described ^21^). Each ectopic peak was reformatted into 50-bp sliding windows (in both orientations, with slide of 1bp). The normalized frequency of each 50-mer candidate windows were evaluated in each ChIP-seq dataset relative to a normalized observed frequency in the corresponding background dataset. Scores were determined as the log transformed normalized value of the ratio between ChIP-seq and background, and those with a score greater than or equal to 2 were included in our study as a high-confident enrichment set.

### Analysis of CENP-A peaks overlap with functional annotation

Ectopic CENP-A peak calls, i.e. those that did not overlap with centromeric α-satellite DNA, were evaluated for enrichment with functional annotation if they were supported between replicate ChIP-seq experiments and overlapped at least one enriched 50mer with a log-transformed normalized ratio >=2, or with a minimum standard ratio of 5-fold. Resulting high-confident ectopic peak calls were intersected (BEDTools: intersect; ^91^ with select functional datasets in the genome (UCSC table browser, ^92^. Peaks that intersect with GRCh38 RefSeq genes (including introns and exons) w/ minimum overlap (-f 0.9;or 90%) required as a fraction of SICER peaks, as well as 1000 bp upstream and downstream (with minimum overlap of 1bp with SICER peak). To evaluate the role of expression, gene annotation was catalogued further based on Intersection with ENCODE HeLa expression data (wgEncodeRegTxnCshlLongRnaSeqHelas3CellPapRawSigPooled) with RefSeq gene annotations (where 22,211 RefSeq Genes (40.5% of total) demonstrated at least >=10 average reads/gene; and highly expressed RefSeq Genes (10,033, or 18.3% of total) RefSeq genes are defined as >=100 average reads/gene). To investigate peak overlap with sites of CTCF enrichment, we intersected peaks with two ENCODE replicate datasets: HeLa-S3.CTCF_Ht1.bed and HeLa-S3.CTCF_Ht2.bed (with minimum overlap of 20 bp). To study the overlap with sites of open chromatin peaks were intersected with ENCODE datasets: HeLa-S3.UW_DNase1_HS.Ht1.bed and HeLa-S3.UW_DNase1_HS.Ht2.bed with minimum overlap of 100 bases. Results were evaluated relative to a simulated peak dataset to test if observed peak counts are higher than expected by chance. Simulations were repeated 100x to provide basic summary statisics: average, standard deviation, max/min, relative enrichment value and empirical p-value.

### Repli-seq experiments

BrdU labeled DNA across S phase was prepared as previously described ^56^ with some modifications. Briefly, cells were synchronized using double thymidine block and release ^8^. Following release from double thymidine block, cells were labeled with BrdU (Sigma, B5002) for 1 hour by adding BrdU to the culture medium to a final concentration of 50 µM. For labeling at early-S (S1), BrdU was added immediately after release (S0). For labeling at mid-S (S4), BrdU was added 3 hours after release (S3). For labeling at late-S (S7), BrdU was added 6 hours after release (S6). Following labeling with BrdU, genomic DNA was extracted, sonicated and heat denaturated as previously described ^56^. BrdU labeled DNA was immunoprecipitated using an anti-BrdU antibody (Becton-Dickinson Biosciences, 555627) coupled to magnetic Dyna M-270 epoxy beads (Thermo Fisher Scientific, 14301). Eluted single-stranded DNA was made into double stranded DNA using random-prime extension (Thermo Fisher Scientific, Random Primers DNA Labeling Kit, 18187–013). Following cleanup of the double stranded DNA (QIAgen QiaQuick PCR Purification Kit, 28104), the DNA was validated by performing quantitative real-time PCR using primers for MRGPRE, MMP15, HBE1 and Sat2. Libraries were then prepared as described above, and sequenced using the Illumina instrument per manufacturer’s instructions (Illumina, San Diego, CA) with the exception that following adapter ligation, repli-seq libraries were size selected between 250–500bp.

### Mass spectrometry identification of proteins associating with CENP-A^TAP^ chromatin

CENP-A^TAP^ was immunoprecipitated from the chromatin fraction of randomly cycling cells or late S synchronized cells as described above. Following beads washes, beads were snap frozen in liquid nitrogen. Samples were diluted using 100 mM Tris pH 8.5 to a final concentration of 2 M urea and digested with trypsin (Promega) overnight at 37 degrees Celsius. The protein digests were pressure-loaded onto 250 micron i.d. fused silica capillary (Polymicro Technologies) columns with a Kasil frit packed with 3 cm of 5 micron Partisphere strong cation exchange (SCX) resin (Whatman) and 3 cm of 5 micron C18 resin (Phenomenex). After desalting, each bi-phasic column was connected to a 100 micron i.d. fused silica capillary (Polymicro Technologies) analytical column with a 5 micron pulled-tip, packed with 10 cm of 5 micron C18 resin (Phenomenex). Each MudPIT column were placed inline with an 1200 quaternary HPLC pump (Agilent Technologies) and the eluted peptides were electrosprayed directly into an LTQ Orbitrap Velos mass spectrometer (Thermo Scientific). The buffer solutions used were 5% acetonitrile/0.1% formic acid (buffer A), 80% acetonitrile/0.1% formic acid (buffer B) and 500 mM ammonium acetate/5% acetonitrile/0.1% formic acid (buffer C). A ten-step MudPIT, each step consisting of a 120 minute elution gradient, was run with salt pulses of 0%, 10%, 20%, 30%, 40%, 50%, 60%, 70% and 100% buffer C and 90% buffer C/10% buffer B. The MS/MS cycle consisted of one full scan mass spectrum (400–1600 m/z) at 60 K resolution followed by five data-dependent collision induced dissocation (CID) MS/MS spectra. Charge state exclusion was enabled with +1 and unassigned charge states rejected for fragmentation. Application of mass spectrometer scan functions and HPLC solvent gradients were controlled by the Xcalibur data system (Thermo Scientific). MS/MS spectra were extracted using RawXtract (version 1.9.9) ^94^. MS/MS spectra were searched with the ProLuCID algorithm ^95^ against a human UniProt protein database downloaded on 03–25–2014 that had been supplemented with common contaminants and concatenated to a decoy database in which the sequence for each entry in the original database was reversed ^96^. The ProLuCID search was performed using full enzyme specificity (cleavage C-terminal to Arg or Lys residue). The data was searched using a precursor mass tolerance of 50 ppm and a fragment ion mass tolerance of 600 ppm. The ProLuCID search results were assembled and filtered using the DTASelect (version 2.0) algorithm ^97^. DTASelect assesses the validity of peptide-spectra matches using the cross-correlation score (XCorr) and normalized difference in cross-correlation scores (deltaCN). The search results are grouped by charge state and tryptic status and each sub-group is analyzed by discriminant analysis based on a non-parametric fit of the distribution of forward and reversed matches. A minimum of two peptides was required for each protein identification. All peptide-spectra matches had less than 10 ppm mass error. The protein false positive rate was below one percent for all experiments.

### Quantification and statistical analysis

For all experiments shown, n is indicated in the figure legends. Values represent the mean ± s.e.m (as indicated in the figure legends).

## Data availability

The datasets generated during the current study were deposited at GEO under primary accession number GSE111381.

The datasets generated during the current study were compared to the following publicly available datasets: ENCODE HeLa DNase-seq (DNase-1 hyper sensitive sites; GEO accsession: GSE90432), ENCODE HeLa CTCF ChIP-sequencing (GEO accession: GSM749729 and GSM749739), ENCODE HeLa H3K27me3 ChIP-sequencing (GEO accession: GSM945208), ENCODE HeLa expression data (UCSC Accession: wgEncodeEH000130), ENCODE HeLa-S3 Repli-seq (GEO accession: GSM923449).

## Author Contributions

Y.N-A. and D.W.C. conceived and designed experiments and wrote the manuscript. Y.N-A performed experiments. KH.M analyzed the sequencing data. M.A.M. and O.S. analyzed data and performed experiments. D.F. suggested experiments and provided key experimental input. A.Y.L and B.R prepared sequencing libraries and provided resources. A.A and J.YIII performed mass spectrometry experiments and provided resources.

## Acknowledgments

The authors would like to thank A. Desai, P. Ly and C. Eissler for critical discussion and helpful suggestions, L.E.T Jansen (Gulbenkian Institute, Oeiras, Portugal) for providing reagents. This work was supported by grants (R01 GM-074150 and R35 GM-122476) from the National Institutes of Health to D.W.C., who receives salary support from the Ludwig Institute for Cancer Research.

---

**Supplementary Table S1.** Read statistics for ChIP-seq and Repli-seq experiments. Total number of merged paired-end read (one read per merged two paired-ends) generated for each sample in dataset, the number (and percentage) of those that were >=100bp in length and the number (and percentage) of reads mapping to α-satellites. **Table A, B**. Read statistics for each sample in the CENP-A^+/LAP^ dataset (A) and for the combined replicates in the CENP-A^+/LAP^ dataset (D) in each condition. **Table C, D**. Read statistics for each sample in the CENP-A^TAP^ dataset (C) and for the combined replicates in the CENP-A^TAP^ dataset (d) in each condition. **Table E, F**. Read statistics for each sample in the BrdU Repli-seq dataset (E) and for the combined replicates in the BrdU Repli-seq dataset (F) in each condition.

| Table A. ChIP-seq replicates statistics for CENP-A<sup>+LAP</sup> chromatin |  |  |  |  |
| --- | --- | --- | --- | --- |
| Experiment | Repli<br>cate<br>No | Total number of<br>merged paired-end reads<br>(100bp x 2) | Total (%) number<br>of merged reads<br>$\geq 100$bp | No (%) of merged<br>reads mapping to<br>α-satellites |
| CENP-A<sup>LAP</sup><br>G1 | 1 | 37,088,538 | 30,232,099 (81.5%) | 27,671,623 (74.6%) |
|  | 2 | 40,641,911 | 33,438,763 (82.3%) | 30,342,295 (74.6%) |
| CENP-A<sup>LAP</sup><br>G2 | 1 | 39,939,734 | 32,202,885 (80.6%) | 24,209,577 (60.6%) |
|  | 2 | 32,689,317 | 25,933,735 (79.3%) | 19,566,369 (59.9%) |
| CENP-A<sup>LAP</sup><br>G1 Input | 1 | 31,874,876 | 28,835,317 (90.5%) | 952,554 (2.98%) |

| Table B. ChIP-seq combined replicate statistics for CENP-A<sup>+LAP</sup> chromatin |  |  |  |
| --- | --- | --- | --- |
| Experiment | Total number of<br>merged paired-end<br>reads (100bp x 2) | Total (%) number<br>of merged reads<br>$\geq 100$bp | No (%) of merged<br>reads mapping to<br>α-satellites |
| CENP-A<sup>LAP</sup><br>G1 | 77,730,449 | 63,670,862<br>(81.9%) | 58,013,918 (74.6%) |
| CENP-A<sup>LAP</sup><br>G2 | 72,629,051 | 58,136,620<br>(79.9%) | 43,775,946 (60.2%) |

| Table C. ChIP-seq replicates statistics for CENP-A<sup>TAP</sup> chromatin |  |  |  |  |
| --- | --- | --- | --- | --- |
| Experiment | Replicate No | Total number of merged paired-end reads (100bp x 2) | Total (%) number of merged reads $\geq 100$bp | No (%) of merged reads mapping to α-satellites |
| CENP-A<sup>TAP</sup> RC | 1 | 9,436,346 | 8,464,391 (89.7%) | 4,588,228 (48.6%) |
|  | 2 | 69,039,423 | 54,952,932 (79.6%) | 31,053,433 (45%) |
| CENP-A<sup>TAP</sup> G1 | 1 | 68,776,382 | 54,522,683 (79.3%) | 28,023,214 (40.7%) |
|  | 2 | 51,746,426 | 40,328,176 (77.9%) | 21,534,570 (41.6%) |
| CENP-A<sup>TAP</sup> mid S | 1 | 65,077,481 | 50,476,007 (77.6%) | 33,879,664 (52.1%) |
|  | 2 | 62,298,174 | 48,772,959 (78.3%) | 31,783,490 (51.0%) |
| CENP-A<sup>TAP</sup> G2 | 1 | 49,206,313 | 40,088,665 (81.5%) | 25,204,739 (51.2%) |
|  | 2 | 60,985,667 | 48,764,504 (80.0%) | 33,414,704 (54.8%) |
| H3.1<sup>TAP</sup> RC | 1 | 31,819,170 | 27,823,087 (87.4%) | 668,075 (2.1%) |
|  | 2 | 42,069,089 | 36,187,314 (86.0%) | 911,027 (2.2%) |
| CENP-A<sup>TAP</sup> RC Input | 1 | 61,557,119 | 61,612,349 (84.5%) | 1,374,644 (2.2%) |

| Table D. ChIP-seq combined replicate statistics for CENP-A<sup>TAP</sup> chromatin |  |  |  |
| --- | --- | --- | --- |
| Experiment | Total number of merged paired-end reads (100bp x 2) | Total (%) number of merged reads $\geq 100$bp | No (%) of merged reads mapping to α-satellites |
| CENP-A<sup>TAP</sup> RC | 78,475,769 | 63,417,323 (80.8%) | 35,641,661 (46.8%) |
| CENP-A<sup>TAP</sup> G1 | 120,522,808 | 94,850,859 (78.7%) | 49,557,784 (41.2%) |
| CENP-A<sup>TAP</sup> mid S | 127,375,655 | 99,248,966 (77.9%) | 65,663,154 (51.5%) |
| CENP-A<sup>TAP</sup> G2 | 110,191,980 | 88,853,169 (80.6%) | 58,619,443 (53%) |
| H3.1<sup>TAP</sup> RC | 73,888,259 | 64,010,401 (86.6%) | 1,579,102 (2.15%) |

| Table E. Repli-seq replicates statistics for BrdU labeled DNA |  |  |  |  |
| --- | --- | --- | --- | --- |
| Experiment | Replicate No | Total number of merged paired-end reads (100bp x 2) | Total (%) number of merged reads $\geq 100$bp | No (%) of merged reads mapping to α-satellites |
| BrdU S1<br>Early S phase | 1 | 48,838,225 | 42,840,706 (87.7%) | 144,337 (0.29%) |
|  | 2 | 938,746 | 730,915 (77.8%) | 2,082 (0.22%) |
| BrdU S4<br>Mid S phase | 1 | 46,828,553 | 40,991,823 (87.5%) | 371,793 (0.79%) |
|  | 2 | 34,480,560 | 31,507,686 (91.4%) | 254,159 (0.73%) |
| BrdU S7<br>Late S phase | 1 | 40,899,839 | 35,827,676 (87.6%) | 1,657,902 (4.05%) |
|  | 2 | 41750,126 | 35,667,382 (85.4%) | 1,682,353 (4.03%) |
| BrdU S1 Input | 1 | 22,887,332 | 21,177,083 (92.5%) | 1,185,850 (5.2%) |
| BrdU S4 Input | 1 | 25,806,345 | 23,810,449 (92.2%) | 1,146,797 (4.4%) |
| BrdU S7 Input | 1 | 25,004,047 | 23,205,784 (92.8%) | 1,322,554 (5.3%) |

| Table F. Repli-seq combined replicate statistics for BrdU labeled DNA |  |  |  |
| --- | --- | --- | --- |
| Experiment | Total number of merged paired-end reads (100bp x 2) | Total (%) number of merged reads $\geq 100$bp | No (%) of merged reads mapping to α-satellites |
| BrdU S1<br>Early S phase | 49,776,971 | 43,571,621 (82.8%) | 146,419 (0.25%) |
| BrdU S4<br>Mid S phase | 81,309,113 | 72,499,509 (89.4%) | 625,952 (0.76%) |
| BrdU S7<br>Late S phase | 82,649,965 | 71,495,058 (86.5%) | 3,340,255 (4.04%) |

**Table S2.**
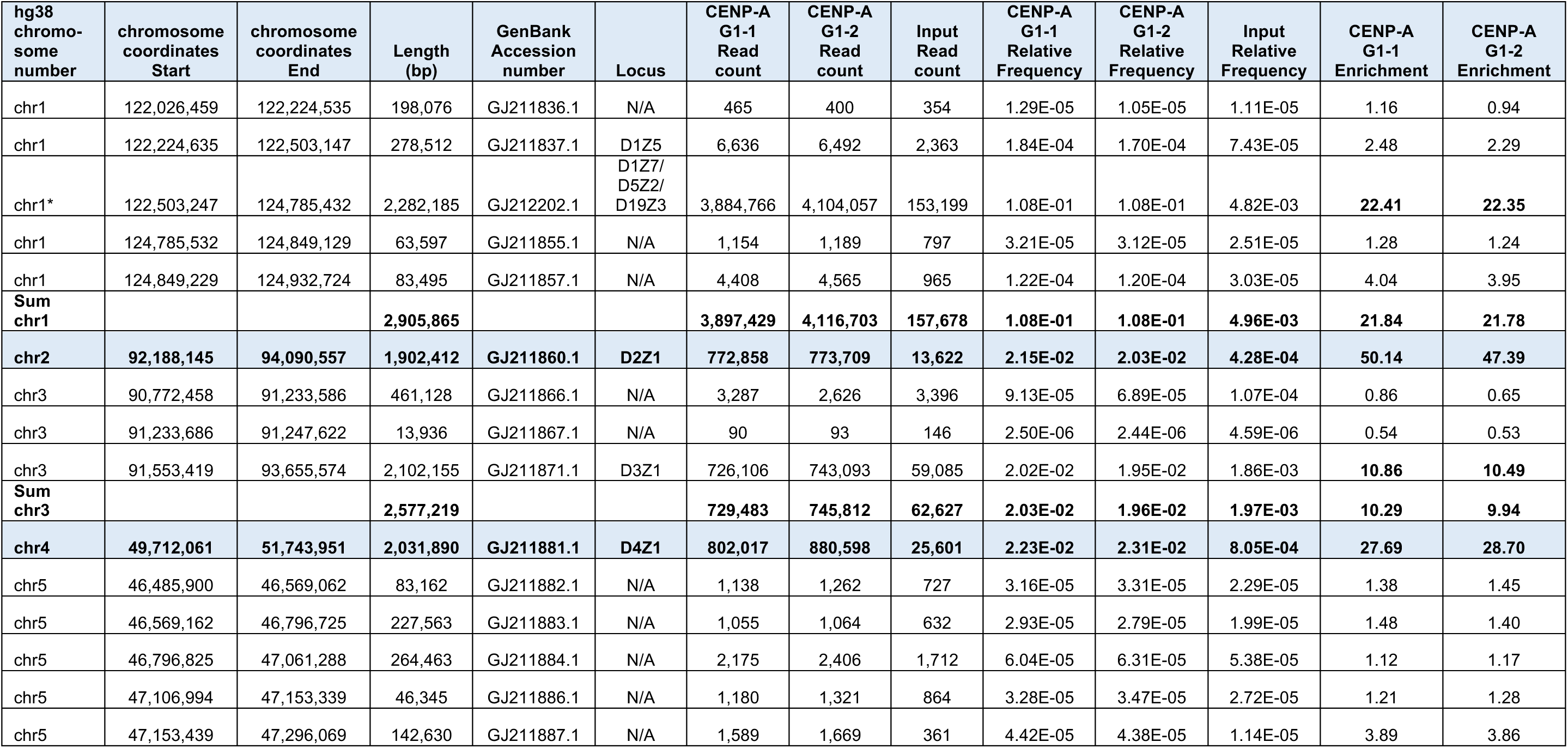
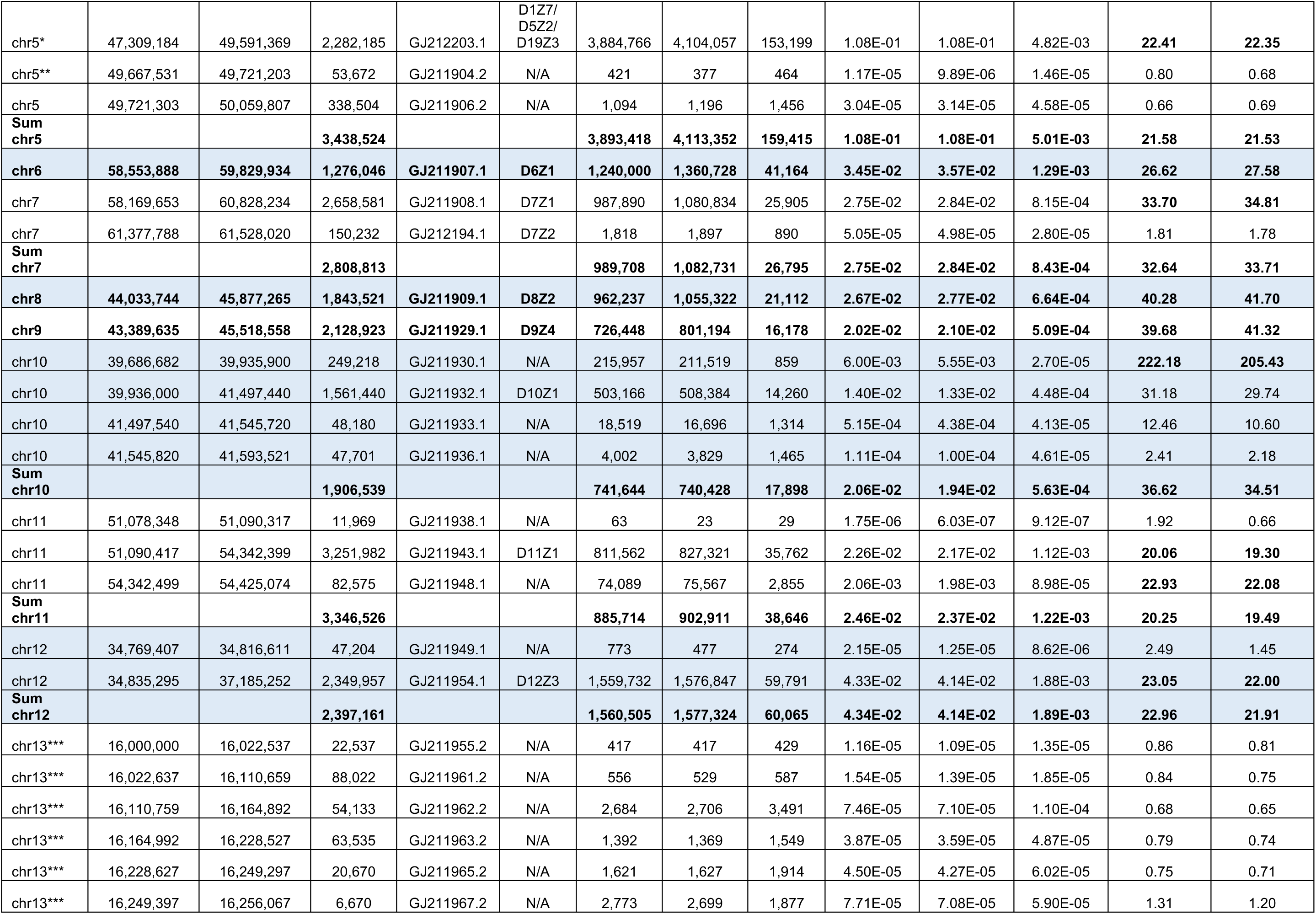
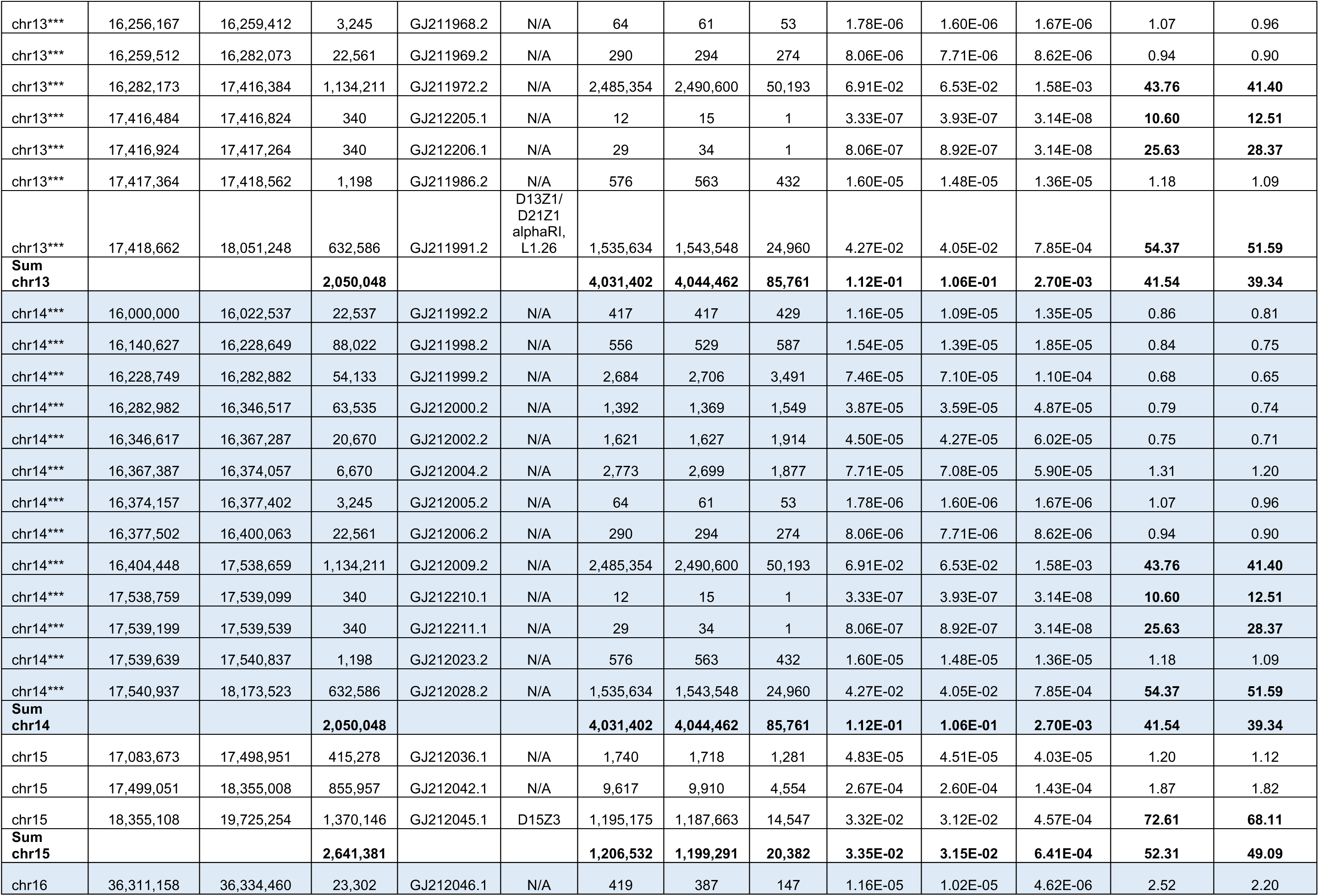
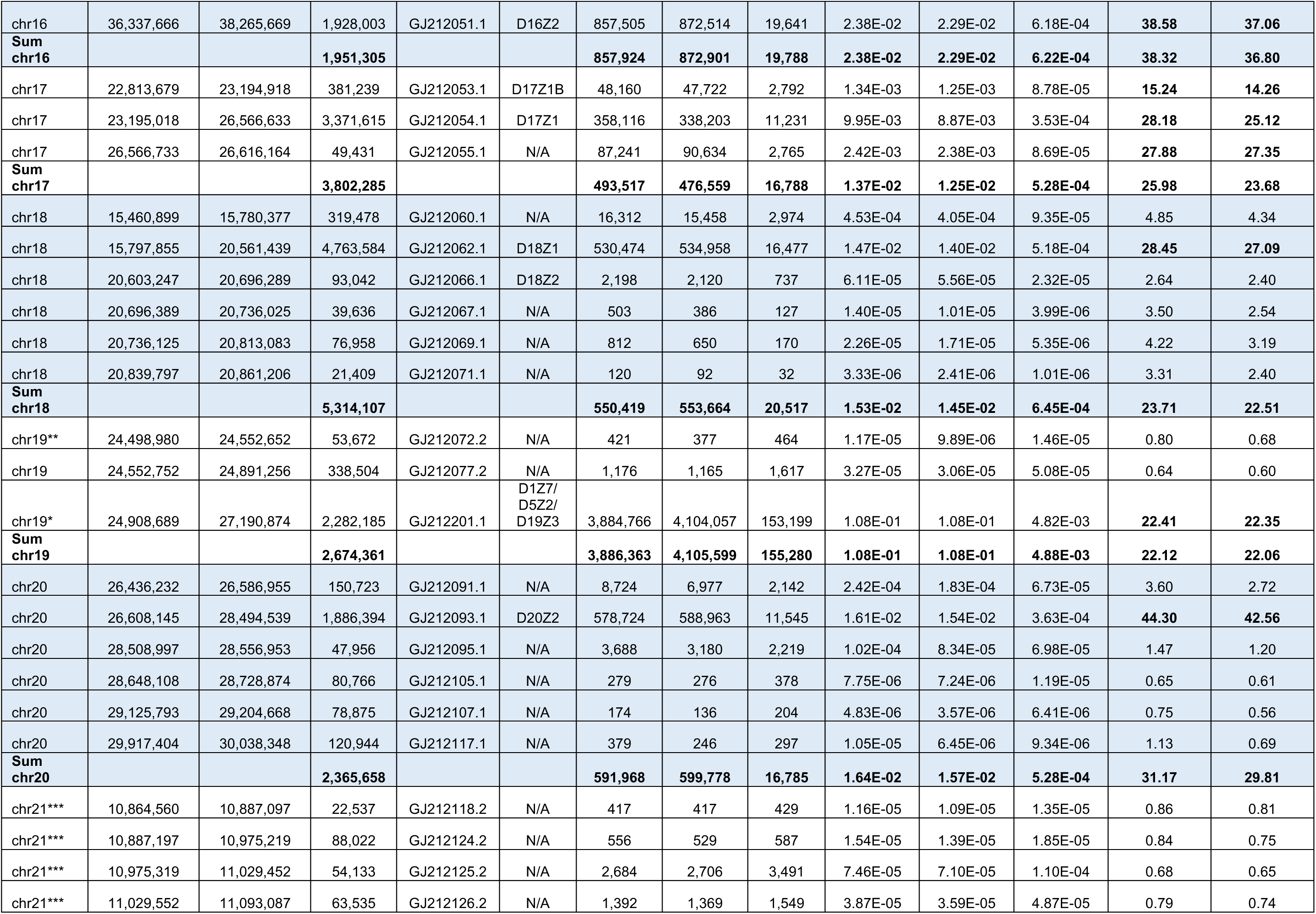
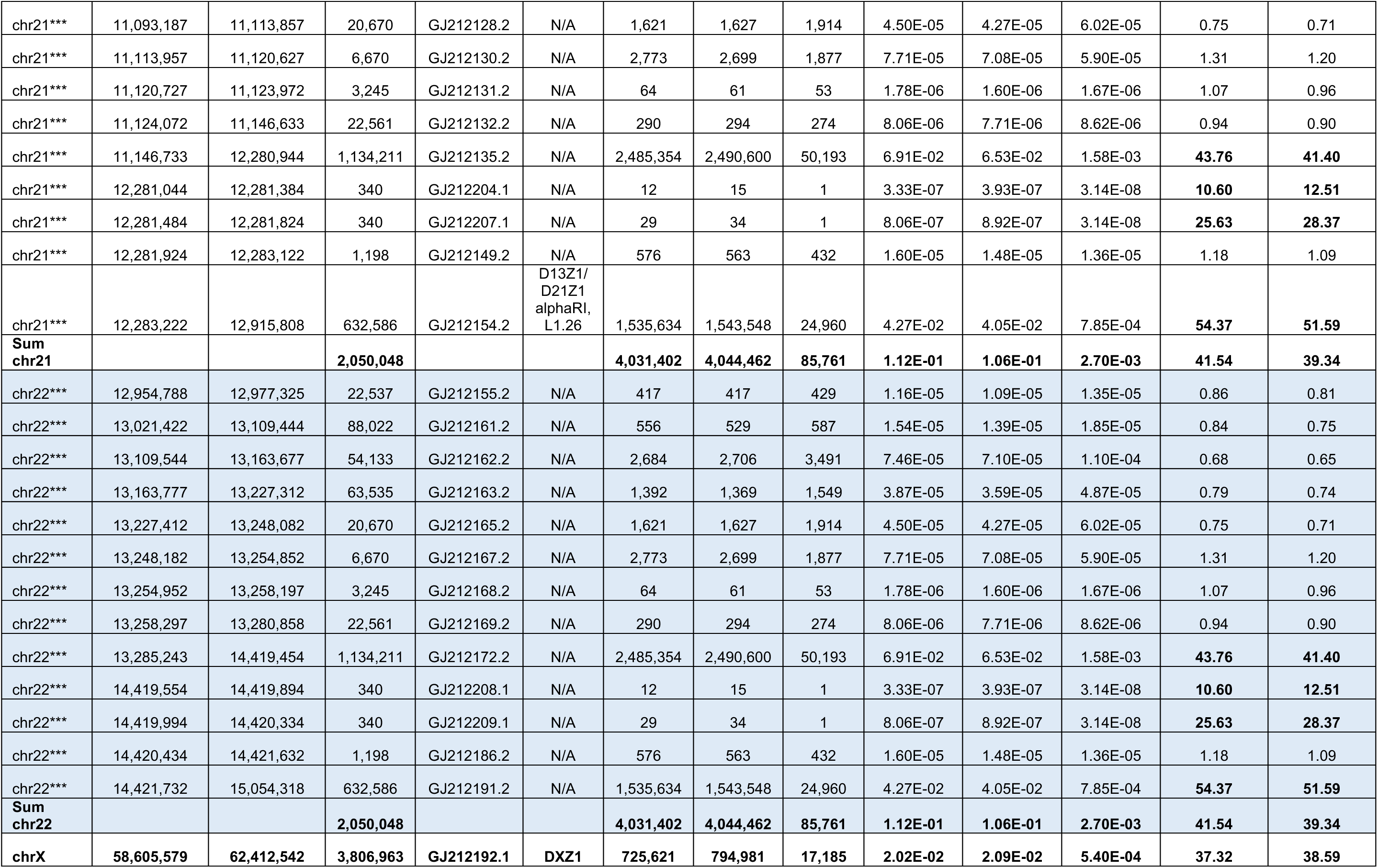
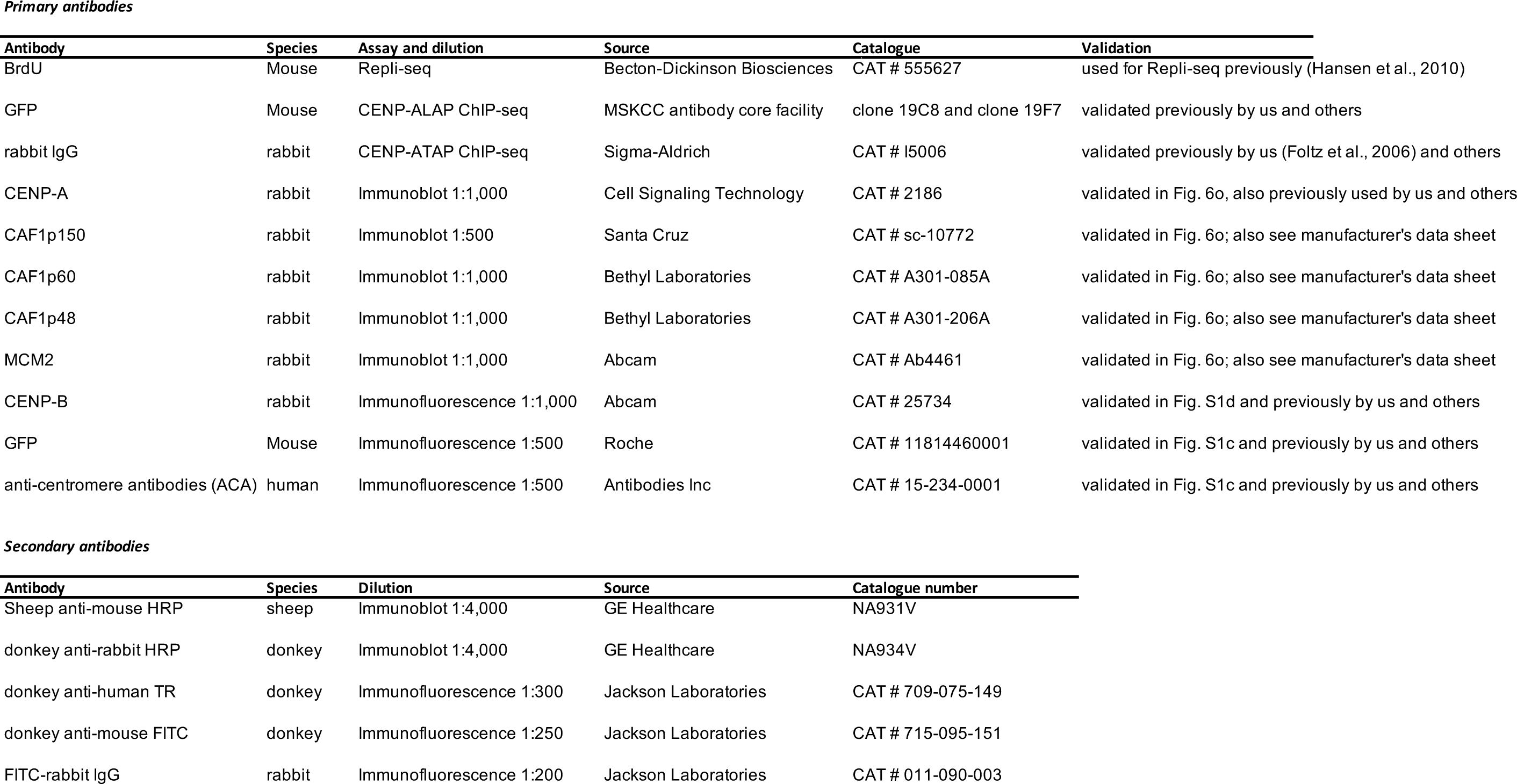
Endogenous CENP-A sequence mapping onto α-satellite DNAs in human centromere reference models for each autosome and the X chromosome. Centromere reference models are from Miga et al. (^46^, unpublished), generated with methods as previously described ^45^. Length estimates are expected to be averaged across arrays from homologous chromosomes. **Column 1:** chromosome information, **column 2:** chromosome start position, **column 3:** chromosome end position, **column 4:** length in bp of each reference model as represented in the human assembly ^46, 8^, **column 5:** Genbank accession, **columns 6:** Genomic locus, if applicable, **column 7, 8, 9:** number of reads for CENP-A^LAP^ G1, replicate samples 1 and 2, and input, respectively, that aligned to the α-satellite reference model, **columns 10, 11, 12:** relative frequency of alignment to the α-satellite reference model is given for CENP- A^LAP^ G1, replicate samples 1 and 2, and input, respectively. **Columns 13, 14:** fold-enrichment of CENP-A^LAP^ G1, replicate samples 1 and 2 at the α-satellite reference model, relative to input. A summary of the reads and bases is given for those chromosomes that have several α-satellite reference models. Arrays that are identical between different chromosome locations are indicated as follows: ^*^Sum of three near-identical arrays on chr1, 5, and 19; ^**^Sum of two near-identical arrays on chr5, and 19; ^***^Sum of acrocentric near-identical arrays on chr13, 14, 21 and 22. Sequence coordinates refer to the human GRCh38 assembly.

